# Evolutionary plasticity of bacterial surface layer protein exoskeletons

**DOI:** 10.1101/2025.04.02.646754

**Authors:** Anna Barwinska-Sendra, Paula S. Salgado, Kacper M. Sendra

**Affiliations:** Biosciences Institute, Faculty of Medical Sciences, Newcastle University, Newcastle upon Tyne, UK

## Abstract

All cellular life possesses environmental interfaces like cell membranes or cell walls, yet the compositional complexity of these major cell components limits structural and evolutionary studies. Prokaryotic surface layer (S-layer) exoskeletons, with their homopolymer paracrystalline architecture, offer a more tractable evolutionary model of an environmental interface. In this study, we reveal the functional, structural and evolutionary diversity of S-layers in Gram-positive *Peptostreptococcaceae*, including pathogens and cancer-promoting species. We uncover novel S-layer architectures with diverse biochemical and physiological properties, enabled by a modular design co-evolving with other cell envelope components. We elucidate the mechanisms and evolutionary pathways underpinning the emergence of novel S-layers and the diversification of existing ones. Our findings establish the S-layer as a paradigm of cellular and molecular modularity and evolutionary plasticity. The demonstrated adaptability of these biological exoskeletons enables rapid reconfiguration of bacterial cell surface architecture and physiology, facilitating immune evasion in pathogens.

## Introduction

The cell wall and cell membrane usually constitute the core components of the bacterial cell envelope, the interface between a bacterial cell and its environment. The layered architecture of the cell envelope can be further augmented by the presence of an outer cell surface-layer (S-layer) exoskeleton. S-layers are found across Archaea, Gram-positive, and Gram-negative Bacteria(*1–4*), are visible using high-resolution imaging techniques, and their building blocks are some of the most abundant cellular proteins. The S-layers studied so far are generally composed of a major protein that assembles into the paracrystalline lattice, and S-layer-associated proteins that confer additional functionality(*5*), as observed in pathogens such as *Bacillus anthracis*(*6*) and *Clostridioides difficile*(*7*). Proposed physiological roles of S-layers and S-layer proteins include protection against cell wall degradation and osmotic stress(*8–11*), biofilm formation(*12–14*), adhesion(*15–17*), phage recognition(*18, 19*), and/or function as a molecular sieve(*20, 21*). Understanding the biology of S-layers is significant within biotechnology, with potential applications as scaffolds in bio-product design(*22, 23*), drug nanocarrier(*24*) or vaccine delivery system(*25*), and within healthcare, since they have been implicated in immunomodulation by microbiota important for optimal health(*26–29*) and in the infection process of different pathogens(*30–32*), and are a target for the development of novel therapeutics(*32, 33*).

Although the diversity of S-layers across different, distantly related bacterial taxa is well established(*2, 34, 35*), general evolutionary mechanisms shaping this diversity across the Tree of Life remain unexplored. S-layer proteins, including those sharing the same structural fold, often lack any identifiable sequence homology(*2*). This complicates studies of S-layer origins and evolution and provides a major challenge for the identification of S-layer proteins and prediction of conserved S-layer functions such as protection against the human immune system. Recent accomplishments in structure-based predictive algorithms provide major advancements in our capacity to predict S-layer protein candidates and S-layer architectures(*34, 35*). However, due to their extreme sequence variation and low number of experimentally verified and studied S-layers, our understanding of functional diversity, ecological significance, and evolution of this important element of prokaryotic cell biology is still limited.

The evolutionary origins of these paracrystalline layers surrounding entire prokaryotic cells, including that of many pathogens, are also not clear. It is commonly hypothesised that S-layers have originated multiple independent times(*34, 36*), mainly based on the sequence and structural diversity of S-layer proteins and lattices(*34*). However, the observed variation of surface exoskeletons could, at least partially, also be explained by an alternative hypothesis of a common origin followed by an extreme divergence and lateral transfer of S-layer proteins. In pathogenic *C. difficile*, the S-layer protein (*Cd*SlpA), encoded within the genomic S-layer cassette, is shared across *C. difficile* isolates *via* homologous recombination(*37*), and its highly variable sequence enables its use for typing (S-layer cassette type, SLCT) of different clinical isolates(*37, 38*). A homologue of *Cd*SlpA, *Peptostreptococcus anaerobius* surface protein PCWBR2 was recently implicated in promoting colorectal carcinogenesis(*39*). Despite their medical significance, our understanding of the structural and functional landscape of S-layers formed by these diverse orthologues from closely related species is limited and current experimental structural information is available for only two *C. difficile* sequence types(*40*).

Here, to address these major gaps in our understanding of S-layer biology and to circumvent the inherent challenges in studying these cellular structures, we focus on narrower phylogenetic scales, where S-layer evolution is more tractable. We experimentally characterise novel S-layer types and present evidence of independent S-layer losses and gains across a bacterial family *Peptostreptococcaceae*. We reveal the role of modular evolution in shaping S-layer diversity and enabling anchoring to variable cell wall architectures. We present novel structures of *C. difficile* S-layers and uncover how, at the narrowest phylogenetic scales, extreme sequence divergence, recombination, and adaptive evolution drive extensive structural changes of the crystalline lattice surrounding the bacterial cells, without compromising its role in protection against the human immune system. Physiological tests and CRISPRi experiments demonstrate that the resistance to the innate immunity effectors emerged specifically in pathogenic lineages. Our results provide insights into the mechanisms of adaptation of host-exposed cell surfaces of S-layer-containing pathogens, and the evolution of the environmental interfaces across the bacterial Tree of Life.

## Results

### The evolutionary and structural diversity of S-layers across the Tree of Life

It is often hypothesized that S-layers are common in Prokaryotes(*1–3*). To test this hypothesis, we compiled a list of previously studied examples of S-layer-positive species and S-layer proteins with strong experimental evidence identified in the available literature, including verification using high-resolution imaging and molecular techniques (Supplementary Data 1). Mapping this evidence onto the Tree of Life (Fig. 1a) revealed that, although verified S-layer-positive species are distributed across many lineages, they represent only a small subset of the analysed Bacteria and Archaea. This reflects the disparity between the number of known species of Prokaryotes and those that have been cultured(*41*), of which only a small number have been tested for the presence of S-layers. Our analysis highlights the extensive gaps in the current understanding of S-layer diversity and distribution.

**Fig. 1:**
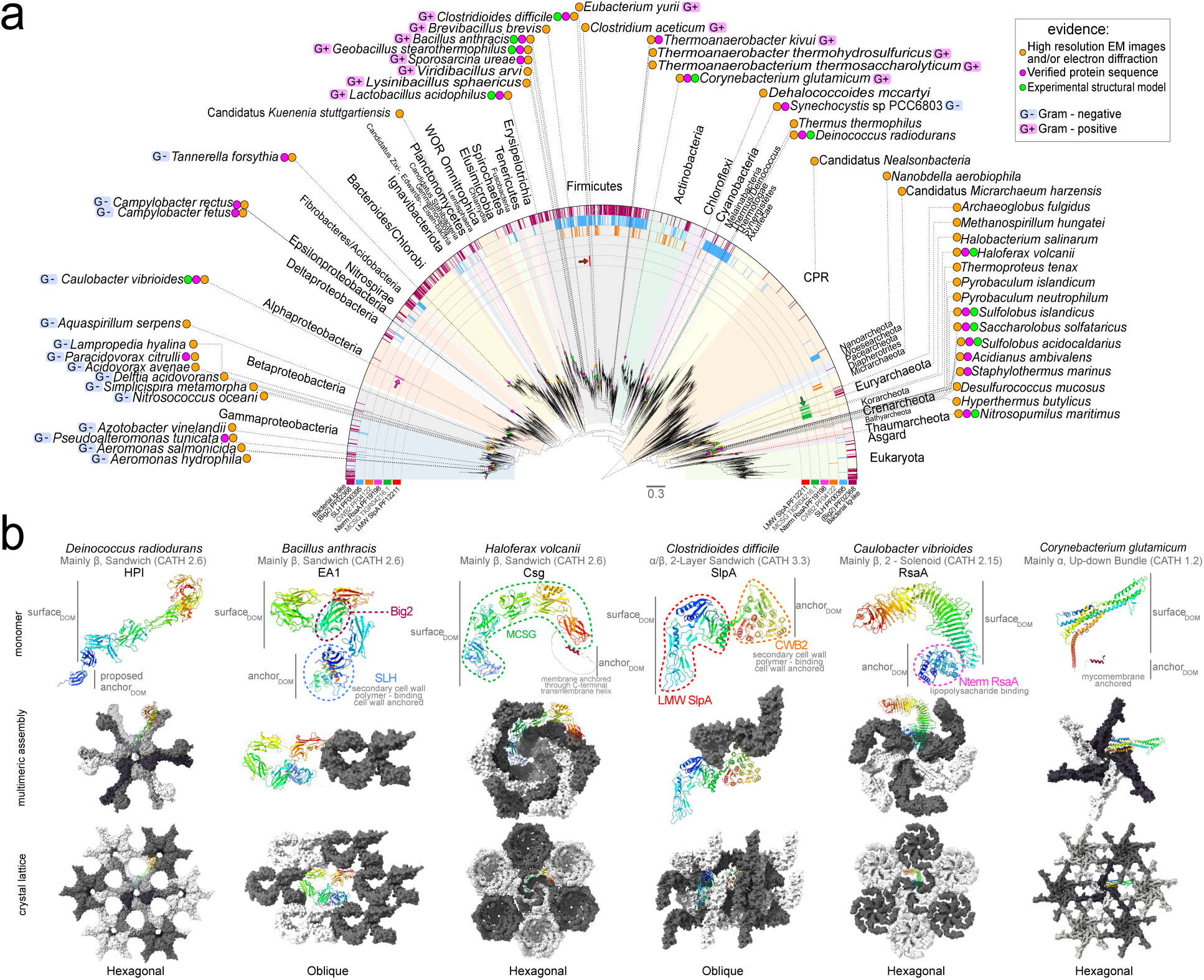
Diversity of S-layer protein lattices across the Tree of Life. **a,** Species names of representative 47 Bacteria and 21 Archaea with previously experimentally verified S-layers (Supplementary Data 1) were annotated onto a phylogeny of 2,705 Bacteria, 315 Archaea, and 149 Eukaryota. Coloured circles represent the available experimental evidence including verification *via* high-resolution imaging techniques (orange), structural biology (green), and protein sequence verification of SLLP (magenta). The coloured semi-circles represent hmmsearch results using HMM profiles of domains found in SLLPs with known structures presented in **b**, demonstrating the presence of both common (e.g. Big2 and SLH), and species- or lineage-specific domain profiles (coloured arrows, e.g. LMW SlpA, SLP_L_). **b,** Cartoon representations of structural models of SLLPs (monomer) and surface representations of their higher order organisation (multimeric assembly and crystal lattice) within the S-layer. The sequence regions (dashed outlines) identified with the available HMM profiles (**a** and Supplementary Fig. 2) can be found within two distinct domains, mainly involved in either cell surface anchoring (anchor_DOM_) or paracrystalline S-layer lattice formation (surface_DOM_). The experimental surface_DOM_ structures (Supplementary Fig. 1a) belong to at least four types of structural folds identified using Foldseek (CATH: 2.6, 3.3, 2.15, 1.2). Multiple monomers (shades of grey) within the multimeric assemblies assume diverse higher-order geometries, which constitute a repeating unit within the crystal lattice with varying degrees of complexity and symmetry types (e.g. hexagonal and oblique). These diverse S-layer architectures are attached to the cell surfaces using a variety of mechanisms including membrane anchoring through transmembrane helices, and binding to different secondary cell wall polymers (SLH and CWB2) or lipopolysaccharides (Nterm RsaA). Structures of the previously determined representative S-layer lattice proteins (8CKA, 8OPR, 7PTR, 7ACY, 6Z7P, 9GK2) were visualised using ChimeraX.

To address these gaps, we investigated the distribution of homologues of the known full-length S-layer protein sequences and their constituent domains across the Tree of Life using protein sequence homology searches (Supplementary Fig. 1), hidden Markov model (HMM) profile searches (Supplementary Fig. 2), and protein similarity network analyses (Supplementary Fig. 3). The main building block of the paracrystalline S-layer array, here referred to as Surface Layer Lattice Protein (SLLP), usually displays a bipartite architecture(*2*). It consists of a cell anchoring domain (anchor_DOM_), which often represents a general anchoring mechanism of cell surface proteins, and an S-layer ‘crystallisation’ or surface-forming domain (surface_DOM_), which plays a more specific role in forming the paracrystalline lattice layer exposed to the extracellular environment (Fig. 1b and Supplementary Fig. 1a). Anchoring domains such as SLH have additional physiological roles, including outer membrane tethering in diderm Negativicutes(*42*), and in our analysis, are found across a broad taxonomic range, often in multiple proteins other than SLLPs from the same genome (Supplementary Fig. 3g). In addition, multiple different types of anchor_DOM_s are often found within a single cell-anchored protein family (Supplementary Fig. 3b, d), indicating that they can be swapped during cell-anchored protein evolution. Therefore, anchor_DOM_s are not specific to SLLPs and constitute swappable protein modules that enable the binding of surface proteins to alternative cell envelope architectures. This likely reflects the structural co-evolution of different cell-envelope layers, such as the cell wall and the S-layer.

In contrast to the anchor_DOM_s, specific surface domain sequence profiles are less common and display narrower taxonomic distribution, suggesting lineage-specific adaptations and more specific functions in S-layer formation (Fig. 1a and Supplementary Fig. 3c). The majority of the analysed surface_DOM_s, including those sharing the common immunoglobulin-like domain architecture (Fig. 1b, and Supplementary Fig. 1a), display extensive sequence diversity. The substantial discontinuity in the structural and sequence landscape of bacterial S-layers is further apparent in SLLPs sharing similar domain folds (Fig. 1b, and Supplementary Fig. 1a) but no sequence homology (Supplementary Figs. 1c, 2a, 3b) like bacterial *B. anthracis* EA1 and *Deinococcus radiodurans* HPI, or archaeal *Haloferax volcanii* Csg, which assemble into extremely diverse paracrystalline layers (Fig. 1b) and utilise different cell surface anchoring mechanisms (Fig. 1b, and Supplementary Fig. 1a).

Combined with the identified gaps in the appreciation of S-layer diversity and distribution, and the possible existence of SLLPs with distinct surface_DOM_ folds, such as *Tannerella forsythia* TfsA (Supplementary Fig. 1a), our analyses are consistent with the existence of undiscovered diversity of prokaryotic exoskeletons in nature. However, the lack of sequence homology often observed between SLLPs belonging to the same structural family and the utilisation of common structural folds complicate analyses of their evolution over wide phylogenetic scales. Given the structural and sequence diversity of surface-forming domains and the multitude of anchoring mechanisms, it is likely that the various S-layers have either emerged multiple independent times from non-SLLPs and/or are a result of extreme surface_DOM_ divergence and anchor_DOM_ swapping among existing S-layer proteins.

### Novel SLLPs form structurally diverse S-layers within a single family of Bacteria

To experimentally investigate the diversity of the S-layer exoskeletons at narrower phylogenetic scales, we first identified the lineages with a high degree of variation in the known SLLP domains. Our analyses across the Tree of Life (Fig. 1a) revealed an interspersed distribution of different anchoring domains – SLH and CWB2 – in Firmicutes. This bacterial phylum includes two important pathogens*, Bacillus anthracis* and *Clostridioides difficile*, with two distinct S-layer surface domain folds (Fig. 1b). Within Firmicutes, *Peptostreptococcaceae* represent a particular ‘hotspot’ of SLH/CWB2 anchoring domain repertoire variation. Therefore, this lineage was selected for combinatorial bioinformatic, cell biology, microbiological, and structural biology analyses to characterise their S-layers and SLLPs and unravel evolutionary mechanisms shaping their diversity.

The experimental investigations (Fig. 2a and d, and Supplementary Fig. 4a) revealed four *Peptostreptococcaceae* (29%, 4/14 species) with no apparent S-layer in whole cell electron micrographs, no ‘ghosts’ in S-layer electron microscopy (EM) preparations, and no abundant protein bands (Supplementary Fig. 4a), typically observed for SLLPs, in surface protein preparations from S-layer-positive species. In three out of four S-layer-negative species, the identified instances of apparent S-layer absence had scattered distribution across the phylogeny, and no identifiable genomic SLLP homologue sequences, consistent with multiple independent losses of S-layer (Fig. 2d). The existence of multiple S-layer-negative lineages of Bacteria could lend some support to the hypothesis that S-layers have evolved multiple independent times(*2, 34*). However, the presence of life cycle-specific or environmentally induced S-layers, or S-layer loss in laboratory conditions, cannot be fully excluded. This is apparent in previously described difficulties in the detection of S-layers in some bacteria(*43*) and, in our dataset, in the single case of *Romboutsia hominis* RFIFI, which contains an SLLP homologue but displays an S-layer-negative morphotype (Fig. 2a and d, and Supplementary Fig. 4). Therefore, currently available data suggests the possibility of independent S-layer losses within *Peptostreptococcaceae*, although these represented the minority of the studied species.

**Fig. 2:**
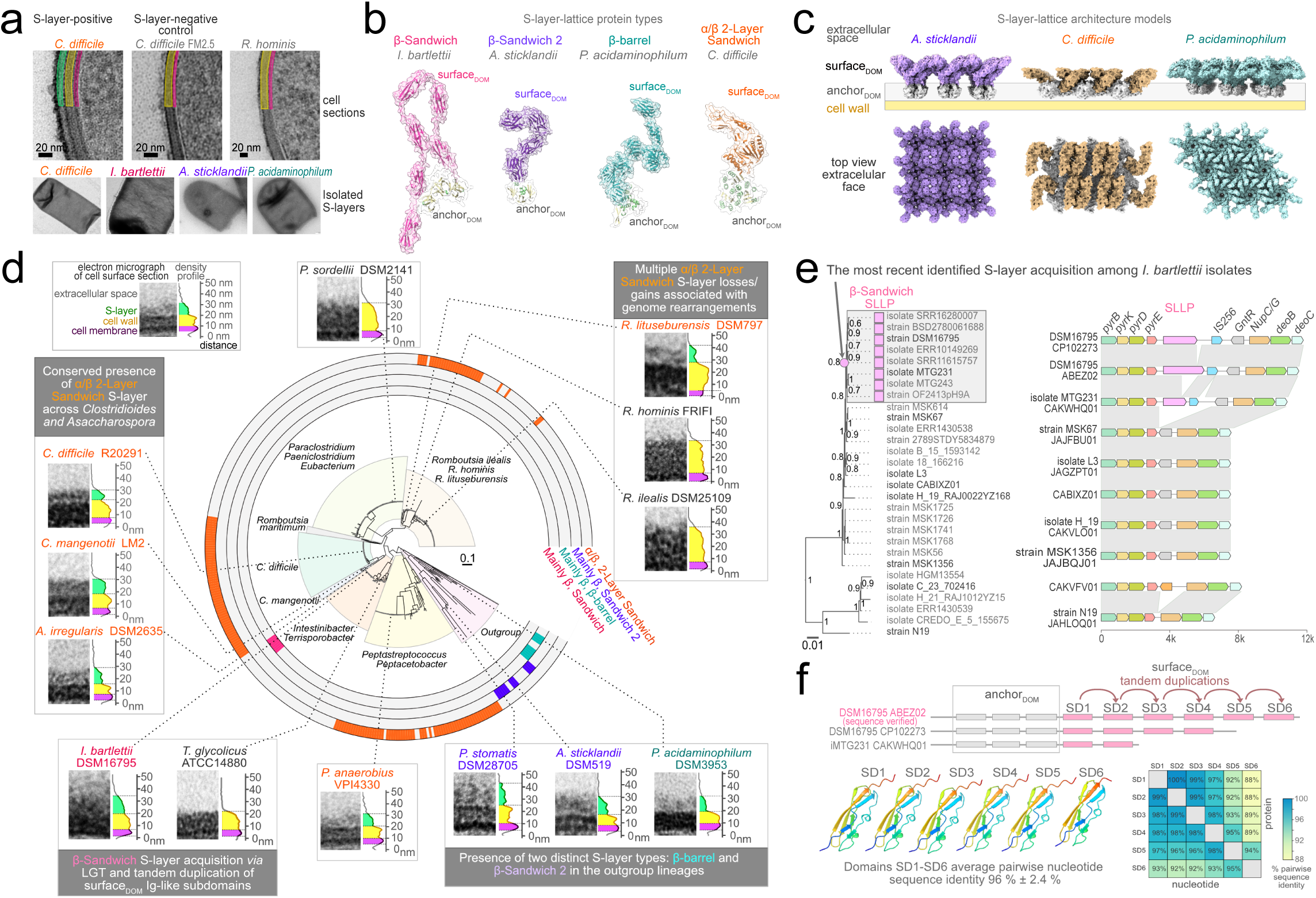
Identification of new types of S-layer and mechanism of emergence of novel S-layer protein. **a,** Representative electron micrographs of uranyl acetate and ruthenium red stained vegetative cells (top panel) of S-layer-positive *C. difficile,* S-layer-negative control of *C. difficile* FM2.5, and *R. hominis* with no apparent S-layer. The coloured regions correspond to S-layer (green), cell wall (yellow), and cell membrane (pink). S-layers (bottom panel) can be stripped from the vegetative cells due to the structural stability of their paracrystalline lattice. **b,** Determined crystal structure (*C. difficile)* and AlphaFold2 predicted structural models of the S-layer lattice proteins (SLLPs) identified by mass spectrometry. The experimentally verified *Peptostreptococcaceae* SLLPs belonged to one of the four distinct surface domain (surface_DOM_) folds. **c,** Surface representations of S-layer models, including top-scoring SymProFold predictions (*A. sticklandii, P. acidaminophilum*), demonstrate that different *Peptostreptococcaceae* SLLP surface_DOM_s (coloured) can assemble into distinct lattice architectures on the bacterial cell surface. **d,** Species tree mapping of the *Peptostreptococcaceae* experimentally tested for the presence of S-layers and SLLPs. Density profiles next to the electron micrographs of the cell envelopes were colour-coded to indicate the location of the S-layer (green), cell wall (yellow), and cell membrane (pink). The scale bars measure distance (in nm) from the bottom of the micrograph. Species names were coloured based on the structural classification (**b**) of the verified SLLP surface_DOM_ of *Peptostreptococcaceae* with apparent S-layer or those with no evidence of S-layer (black). Coloured semi-circles represent the distribution of the protein homologues of each SLLP-type identified in 453 *Peptostreptococcaceae* genomes. **e,** The presence of the β-sandwich SLLP (pink rectangles) was mapped onto the species tree of *I. bartlettii* isolates (left panel). The recent acquisition of the SLLP (pink circle) is apparent in genomic region alignment (right panel), indicating ancestral insertion of the SLLP (pink) and IS256 transposase (blue) coding genes within a broadly conserved nucleotide metabolism locus. **f,** Domain architecture of the *I. bartlettii* SLLPs (top panel) demonstrates a possible variation in the number of the β-sandwich surface_DOM_ subdomains (SD1-6) among different isolates. All six SD1-6 (bottom panel, left) share high levels of pairwise protein and nucleotide sequence identity (bottom panel, right) consistent with SLLP origin *via* multiple tandem subdomain duplications.

In the majority (71%, 10/14 species) of the studied *Peptostreptococcaceae*, we found evidence of S-layers and SLLPs (Fig. 2d; and Supplementary Fig. 4a). AlphaFold2 structural predictions and Foldseek analyses revealed that six of the identified SLLPs (60%, 6/10 SLLPs) contained surface domains homologous to the α/β-sandwich structural fold (Fig. 2b, c), previously only found in *C. difficile* SLLP (*Cd*SlpA). This type of SLLP, with only two or three surface-exposed subdomains (Fig. 2c), features a distinctively shorter surface_DOM_ than other identified SLLP types and can potentially assemble into a more compact S-layer lattice, as observed in *C. difficile*. Another distinction of this SLLP type is the higher proportion of α-helices within its surface_DOM_(*40*), as opposed to predominantly β-strand surface_DOM_s of S-layers commonly found across the Tree of Life (Fig. 1b). Four of these shorter SLLPs were found in close relatives of *C. difficile*, including *Asaccharospora irregularis* and *Clostridioides mangenotii*, suggesting that this type of S-layer was present in their last common ancestor. Interestingly, the SLLP from *Peptostreptococcus anaerobius* contained the same fold, and our experiments identified it as PCWBR2 (Supplementary Fig. 4a), a protein implicated in promoting colorectal carcinogenesis(*39*).

Three other identified *Peptostreptococcaceae* SLLPs (30%, 3/10) contained β-sandwich surface_DOM_s, with two from the most basal *Peptostreptococcaceae* outgroup lineages (Fig. 2a-d and Supplementary Fig. 4). Interestingly, while α/β-sandwich surface_DOM_ shared enough sequence homology to group weakly within a single protein similarity network, the β-sandwich fold SLLPs grouped within two distinct networks (Supplementary Fig. 4b). The more broadly distributed β-sandwich S-layers were found within the outgroup lineages including *Acetoanaerobium sticklandii* and *Peptoanaerobacter stomatis*. Their broader distribution across the species tree suggested more ancient S-layer acquisition, contrasted by the second type of β-sandwich S-layers, found only in a narrow subsample of closely related *Intestinibacter bartlettii* genome assemblies (Fig. 2e). Inspection of all available *I. bartlettii* genomes revealed a recent SLLP acquisition likely *via* transposition, indicated by its co-distribution with an adjacent IS256 gene (Fig. 2e) encoding a transposase homologue. Furthermore, structural and sequence comparisons of the *I. bartelettii* SLLP revealed that it has evolved *via* multiple tandem duplications of a single common β-sandwich Immunoglobulin-like (PFAM ID, PF16403) surface subdomain (Fig. 2f). At this stage, we were able to sequence-verify SLLP from the single *I. bartelettii* isolate available in culture collections. The significance of subdomain number variation observed across available *I. bartlettii* genome assemblies remains unclear, as SLLP-coding sequences are often incomplete and found at the end of a contig. Nevertheless, all six surface_DOM_ subdomains of the verified sequence shared, on average, 96 % of pairwise nucleotide sequence identity, consistent with the recent emergence of this β-sandwich SLLP type.

The single SLLP from *Peptoclostridium acidaminophilum* contained the third distinct type of S-layer surface-forming domain, β-barrel (10%, 1/10 SLLPs). Its similarity to the β-sandwich fold suggests that it may have evolved either from a shared β-sandwich SLLP ancestor or originated independently from a non-SLLP β-barrel-containing protein. The β-barrel SLLP contained the largest surface_DOM_ among the identified *Peptrostreptococcaece* SLLPs, with an extended 10-subdomain architecture, previously also found in the β-sandwich S-layer of ammonia-oxidizing marine archaea(*21*). The available crystal structures or calculated SymProFold S-layer lattice models indicate that the β-barrel SLLP can assemble into an hexagonal lattice, in contrast to the oblique lattice determined in *C. difficile* α/β-sandwich S-layer or the predicted square symmetry of the *A. sticklandii* β-sandwich S-layer model (Fig. 2c).

By combining bioinformatics and experimental investigations, we uncovered novel types of SLLPs and S-layers, recent SLLP emergence and S-layer acquisition, and multiple independent SLLP losses in *Peptostreptococcaceae*. The revealed structural diversity of *Peptostreptococcaceae* SLLPs, including multiple distinct S-layer surface-forming folds assembled into multiple distinct S-layer architectures, reflects what we observed in our analyses across the Tree of Life (Fig. 1). Therefore, the presence of diverse S-layer architectures utilising different surface domain folds does not only represent distinct features of distantly related bacterial phyla but can be generated within much shorter evolutionary time frames. The emergence of a novel SLLP-type *via* tandem subdomain duplication in *I. bartelettii* provides an insight into the mechanism likely involved in the origins of novel S-layer architectures.

### Homologous S-layers can impart different resistance to antimicrobial agents

Although the structural and sequence diversity of S-layers at both wider and narrower taxonomic scales is clear, the conservation of their physiological roles remains elusive. Furthermore, the function of S-layers in many organisms has not been experimentally investigated(*2*). S-layers of bacteria such as that of gut pathogen *C. difficile* protect cells from natural antimicrobial agents produced by the host immune system such as lysozyme or human cathelicidin peptide LL-37(*10*). However, it is not known if this is also true for other S-layers we identified across *Peptostreptococcaceae*.

Our initial experimental investigations demonstrated that *Peptostreptococcaceae* S-layers display variable susceptibility to extraction using different chemical methods (Fig. 3a), indicating that at least some of the properties of the outermost cell envelope layer of these bacteria are not conserved. Surprisingly, this variation was not only observed for the S-layers made of different types of SLLPs but also across homologous S-layers formed by the α/β-sandwich SLLPs, including that of *C. difficile* (Fig. 3a). Protein similarity network analyses (Fig. 3b) and manual inspection of multiple sequence alignments revealed extensive sequence divergence within this SLLP-fold. This is particularly apparent in the case of S-layer surface_DOM_s from different isolates of *C. mangenotii* (LM2 and DMS29133), which are not directly connected with each other or with sequences from their closest relatives (*C. difficile* and *A. irregularis*) within the network (Fig. 3b).

**Fig. 3:**
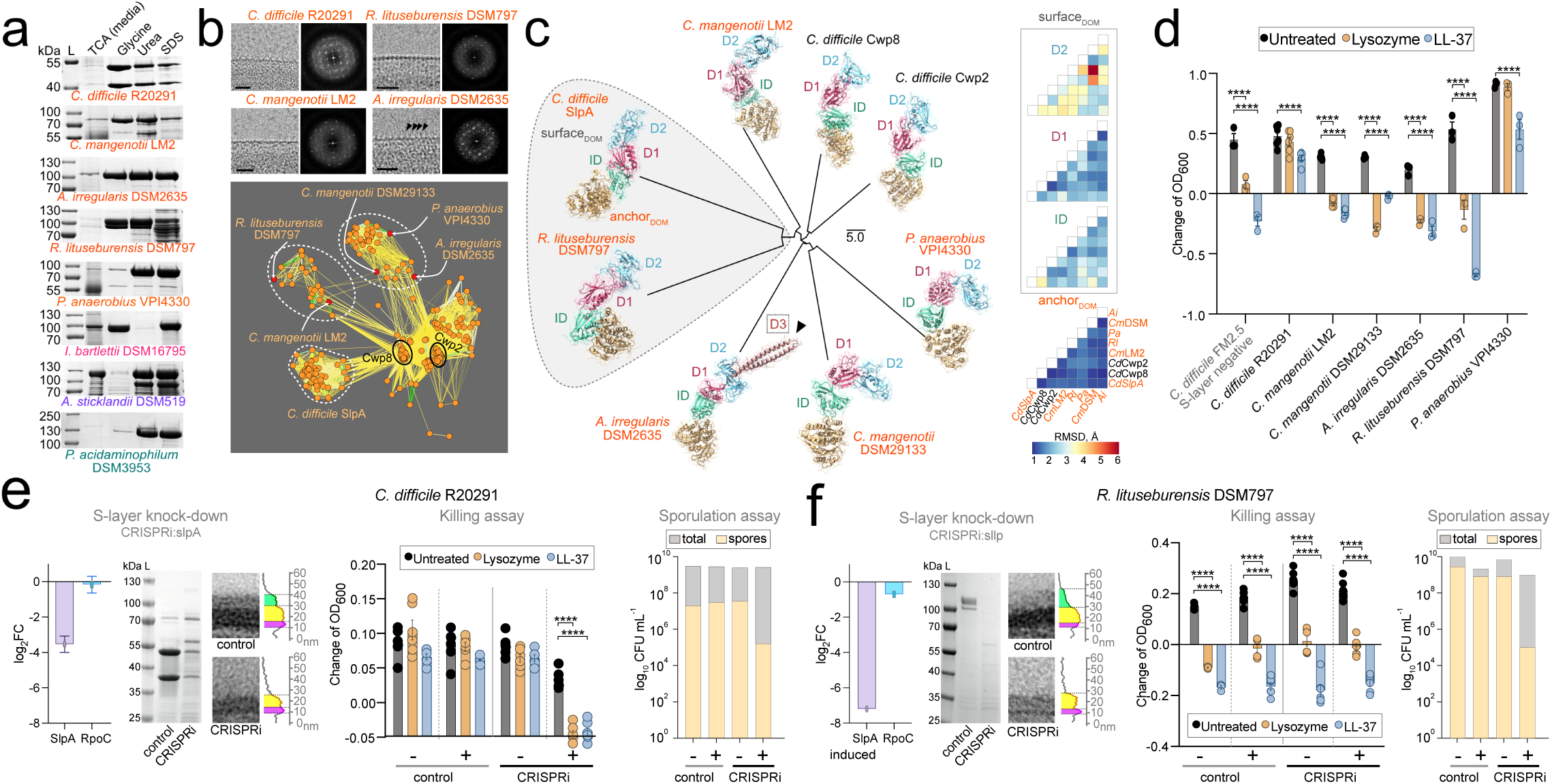
Variable biochemical and physiological properties within a single structural type of S-layer. **a,** SDS-polyacrylamide gels of *Peptostreptococcaceae* low pH glycine, urea, and SDS S-layer preparations. The α/β-sandwich S-layers (orange) displayed contrasting sensitivity to extraction in SDS and glycine. β-sandwich S-layers (pink or purple) varied when extracted in glycine and urea. Trichloroacetic acid (TCA) precipitation determined the SLLP presence in the medium. Molecular weight ladder (L). **b,** Cryo-EM micrographs and Fast Fourier Transform (FFT) diffraction patterns of the α/β-sandwich S-layers (top panel). Additional protrusions were apparent at the extracellular face of *A. irregularis* S-layer (black arrowheads). Protein sequence similarity network (bottom panel) of the α/β-sandwich SLLP surface_DOM_s homologues (orange circles) including S-layer-associated Cwp2 and Cwp8 from the available *Peptostreptococcaceae* genomes. Highly divergent sequences of the verified S-layer surface_DOM_s (red circles) formed three distinct groups (dashed outlines). **c,** The Alphafold3 structural models of identified *Peptostreptococcaceae* α/β-sandwich SLLPs and Cwp2/8 were mapped onto their Dali surface_DOM_s structural comparison dendrogram. Two additional α-helices of *A. irregularis* (dashed rectangle), likely correspond to the S-layer extracellular protrusions (**b**). Heatmap matrices for RMSD values across subdomains D1, D2, ID and anchor_DOM_ show the highest structural divergence within the D2 (average RMSD 3.7 Å). Conversely, the anchor_DOM_ CWB2 fold is highly conserved (average RMSD 1.2 Å). **d,** Broth-based killing assay in the presence of immune effectors (Lysozyme, orange; LL-37, blue) relative to the non-treated control (black). Resistance to growth impairment and cell death was observed only for two pathogenic S-layer-positive species (*C. difficile* and *P. anaerobius*). **e, f**, *C. difficile* CRISPRi SlpA knock-down (**e**) resulted in the sensitivity to both antimicrobial agents in broth-based killing assay (as in **d**), as observed in the S-layer-negative strain FM2.5 (**d**), consistent with S-layer protecting against the host’s innate immunity. The S-layer knock-down in *R. lituseburensis* (**f**) did not change its natural sensitivity (**d**) to lysozyme and LL-37, yet it impacted sporulation in plate-based assay for both species. The gene knock-down was induced with xylose (+) relative to negative plasmid controls (control). CRISPRi SLLP gene knock-downs were validated with RT-qPCR (left panel), SDS-polyacrylamide gels (middle panel) and electron microscopy (right panel). All experiments were performed in biological triplicate, with a standard error of the mean and statistical significance (****, P<0.0001) shown.

Structural comparisons of the experimentally verified α/β-sandwich SLLPs (Fig. 3c) and homologous S-layer-associated proteins Cwp2 and Cwp8, revealed that, despite their extensive sequence variation, the overall fold architecture remains conserved. The general α/β-sandwich SLLP surface_DOM_ consists of three subdomains, equivalent to *C. difficile* SlpA D1, D2 and ID(*40*) and S-layer associated Cwp8 D1, D2, D3(*44*), which are connected via flexible linkers, providing further structural variability as rotation of the relative orientation of the subdomains leads to varying surfaces (Fig. 3b,c). The AlphaFold3(*45*) predictions for the experimentally verified SLLPs fold with three α/β-sandwich subdomains follows the topology seen in *CdI*SlpA, where D1 leads to the loop-rich D2. Insertion of a β-strand into D1 then links to the third subdomain (ID), which connects to the CWB2 anchor_DOM_. The most structurally conserved subdomain in surface_DOM_ is D1, with an average RMSD of 1.7 Å across all SLLPs, while the least conserved is D2 (average RMSD 3.7 Å). The most distinct modification to the general α/β-sandwich architecture was observed in *A. irregularis* DSM2635 where an additional, elongated α-helical region (∼80 Å long) has been acquired and inserted within D2 (Fig. 3c). These additional helices likely correspond to the protrusions extending out of the lattice towards the extracellular environment observed in the cryo-electron micrographs of the *A. irregularis* S-layer preparations (Fig. 3b). Importantly, acquisition of novel types of surface_DOM_ subdomains provides another mechanism of S-layer lattice evolution, alongside variation in the number of subdomains as identified in *I. bartlettii* (Fig. 2f).

Next, we tested if S-layer dependent resistance to innate immune effectors observed in *C. difficile* is conserved in other *Peptostreptococcaceae* containing α/β-sandwich S-layers. Physiological growth tests in the presence of lysozyme and LL-37 revealed variable resistance to these natural antimicrobials (Fig. 3d). Consistent with previous investigation(*10*), S-layer-positive *C. difficile* R20291, but not S-layer-negative *C. difficile* FM2.5, displayed resistance against LL-37 and lysozyme. Interestingly, similar resistance was also observed in cancer-promoting pathogen(*39*) *P. anaerobius* but not in any of the other α/β-sandwich S-layer *Peptostreptococcaceae*, including closest *C. difficile* relatives *A. irregularis* and *C. mangenotii* (Fig. 3d). The presented data is consistent with resistance to these antimicrobials representing specific and potentially independent adaptations of *C. difficile* and *P. anaerobius* cell envelopes to the gut environment.

To further test the variable resistance of α/β-sandwich S-layers to antimicrobial agents, and to experimentally verify that the identified SLLPs are essential for S-layer formation, we performed CRISPRi gene knock-down experiments in *C. difficile* (Fig. 3e) and *R. lituseburensis* (Fig. 3f), two more distantly related *Peptostreptococcaceae* species but with SLLPs sharing a close structural relationship, according to Dali analyses (Fig. 3c). Consistent with the experiments in the wild type-strains (Fig. 3d), knock-down of the SLLP gene abolishes *C. difficile* resistance to lysozyme and LL-37 (Fig. 3e), as opposed to *R. lituseburensis*, where the presence of SLLP provided no apparent protection against either (Fig. 3f).

Previous experiments have also demonstrated that the presence of S-layer plays a role in *C. difficile* sporulation(*10*). In our experiments, CRISPRi S-layer knock-down strains of *C. difficile* and *R. lituseburensis* exhibited sporulation deficiencies in the tested conditions (Figure 3e, f, Supplementary Table 1), suggesting that S-layer integrity plays a conserved role necessary for sporulation of these two α/β-sandwich S-layer containing *Peptostreptococcaceae*. In addition, both mutants displayed a propensity to aggregation in liquid culture, highlighting the conserved role of their S-layers in its prevention.

In summary, despite their structural conservation, homologous α/β-sandwich S-layers from closely related species display a high degree of biochemical and sequence diversity and differ in their contribution to the protection of the cells they surround against natural antimicrobials lysozyme and LL-37. Despite these differences, they also play conserved roles in cellular processes such as preventing cell aggregation and sporulation, important in bacterial survival and infection. Therefore, not all phenotypic effects associated with the presence of homologous S-layers are conserved. Certain functional features could reflect specific ecological niches, such as the gut environment for *C. difficile* and *P. anaerobius*, while more general life cycle adaptations like sporulation and cell-cell interactions might be under broader selective pressure.

### S-layer anchoring domains belong to a broader system of swappable cell attachment modules

The observed diversity of *Peptostreptococcaceae* S-layer surface-forming domains results in multiple S-layer architectures (Fig. 2b-d) with distinct biochemical properties (Fig. 3a) and variable physiological functions (Fig. 3d-f). In addition to surface_DOM_s, SLLPs also contain various anchoring domains that attach the S-layer to the cell. Our analysis across the Tree of Life demonstrated that anchor_DOM_s are not unique to SLLPs, can be found in multiple other proteins encoded in bacterial genomes, and that alternative anchor_DOM_s can be swapped during cell-anchored protein evolution (Supplementary Fig. 3). In pathogenic Firmicutes, S-layers utilise at least two different cell-wall anchoring domains, CWB2 in *C. difficile* and SLH in *B. anthracis*, binding to different secondary cell wall polymers (SCWPs)(*6, 46, 47*). The capacity to utilise swappable cell wall anchors could enable attachment of S-layers to different cell wall components, enabling lateral transfer of S-layers between bacteria utilising different cell wall architectures and facilitating co-evolution of different cell envelope layers. Therefore, we investigated if the *Peptostreptococcaceae* S-layers utilise variable cell anchoring mechanisms and if their anchor_DOM_s evolve as part of the wider cell surface proteome.

The four types of SLLPs we identified in *Peptostreptococcaceae* (Fig. 2b,c) utilise four distinct cell-anchoring domains (Fig. 4a and Supplementary Fig. 5a,b). All the SLLPs with α/β-sandwich surface _DOM_s, including *C. difficile* SlpA, contained CWB2 anchor_DOM_, while the single β-barrel SLLP contained the SLH anchor_DOM_ (Fig. 3a), also found in β-sandwich surface _DOM_-containing S-layers of *Bacillaceae,* including pathogenic *B. anthracis* (Fig. 1b). However, *Peptostreptococcaceae* S-layers with β-sandwich surface_DOM_s contained one of two distinct anchors: SH3 or Cu-oxidase-like N-terminal domain(*48–50*) (Cu-OX, Fig. 3a). This demonstrated that, within the single bacterial family of *Peptostreptococcaceae,* a variety of anchor_DOM_s are utilised by different S-layer lattice-forming surface_DOM_s, reminiscent of the patterns observed across the Tree of Life (Fig. 1 and Supplementary Fig. 1a).

**Fig. 4:**
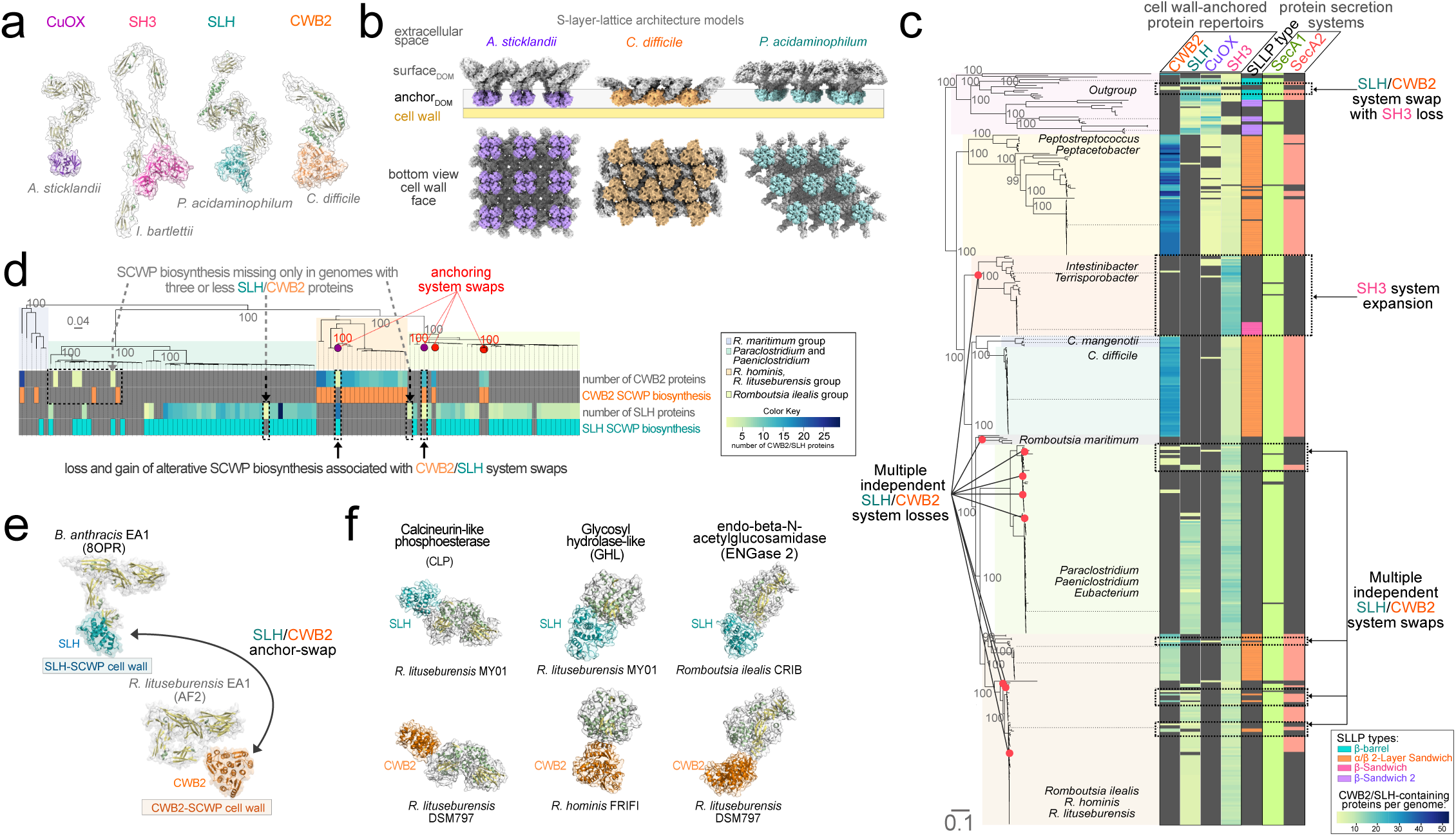
Modular architecture and evolution of the S-layer and other cell envelope systems. **a,** The four experimentally identified *Peptostreptococcaceae* S-layer surface forming fold-types (Fig. 2b) are anchored to the cell using four distinct anchoring domains (orange, CWB2; cyan, SLH; magenta, CuOX; pink, SH3). **b,** Surface representations of S-layer lattice models (as in Fig. 2c) with anchor_DOM_s coloured as in **a**. **c,** Species tree mapping of the total number (yellow-blue heatmap) of cell wall anchored proteins identified in genomes of 453 *Peptostreptococcaceae*. SH3 anchoring is more broadly conserved, while relatively less common CWB2 and SLH systems are interchangeable and were lost (red circles) or swapped (dashed rectangles) multiple independent times. The more general SecA1 (green) secretion system is present in all lineages, whereas SecA2 (red) with an SLLP-specific role in in *C. difficile*, is found primarily with CWB2-anchored S-layers (SLLP type). **d,** Numbers of CWB2 and SLH anchor-containing proteins (yellow-blue heatmaps), and the presence of the secondary cell wall polymer (SCWP) biosynthesis enzymes specific to SLH (cyan) or CWB2 (orange) were mapped onto the *Romboutsia/Paraclostridium/Paeniclostridium* species tree. Enzymes involved in the biosynthesis of SLH- and CWB2-specific SCWPs co-distribute with their respective anchor_DOM_s and are missing only in the genomes containing a low number of anchored proteins (dashed arrows). Multiple independent SLH/CWB2 system swaps (red arrows) and possible complete (red circles) and incomplete (purple circles) system losses (dashed arrows) are apparent. **e,** AlphaFold2 structure predictions of the *R. lituseburensis* homologue of *B. anthracis* SLH-anchored S-layer protein EA1 contains a CWB2 anchor_DOM_ consistent with anchoring domain swap and lateral gene transfer between *Peptostreptococcaceae* and *Bacillaceae*. **f,** AlphaFold2 structure prediction of individual non-SLLP cell wall-anchored proteins such calcineurin-like phosphoesterase (CLP) and peptidoglycan hydrolases (endo-β-*N*-acetylglucosamidase, ENGase; and glycosyl hydrolases, GHL), with swapped SLH/CWB2 anchor_DOM_s.

The utilisation of the four distinct cell wall anchoring domains in *Peptostreptococcaceae* S-layers, including those sharing similar structural folds (Fig. 4a), suggested that these domains may represent swappable modules. Our analyses revealed multiple independent swaps, gains, and/or losses of the entire SLH/CWB2 cell wall anchoring systems within *Peptostreptococcaceae* (Fig. 4c), consistent with the patterns we observed within the broader sampling of Firmicutes (Supplementary Fig. 5). These changes usually affect multiple proteins, as SLH and CWB2 anchor_DOM_ are found on average in 12.5 and 9.8 proteins per anchor_DOM_-containing Firmicute genome, respectively (Supplementary Fig. 5b). Based on protein similarity network analyses, the identified anchoring domain swaps were often observed with retention of the attached functional domain homology (Fig. 4e,f, and Supplementary Fig. 3a-d), suggesting that anchoring system swap does not necessarily reflect changes in the surface proteome functionality. Although at least some of the inferred swaps involved rearrangement of entire genomic regions encoding multiple surface proteins (Supplementary Fig. 6), individual surface proteins are often scattered across the chromosome (Supplementary Fig. 5c). Phylogenetic, structural, and sequence homology analyses are consistent with multiple independent anchor replacements among closely related proteins (Supplementary Fig. 6) indicating that, in addition to the swaps of gene cassettes encoding multiple surface proteins, surface proteomes evolve by replacing anchor_DOM_s in individual proteins. Multiple anchor_DOM_ swaps were also found in species with no evidence of S-layer (Supplementary Fig. 6). Therefore, anchoring domain variation is not specific to SLLPs and S-layers, but it rather reflects a more general trend affecting entire cell wall-anchored protein repertoires.

Neither CWB2 nor SLH systems were found in any of the analysed *Terrisporobacter/Intestinibacter* genomes. Instead, they contained an expanded repertoire of SH3 domain-containing proteins compared to other *Peptostreptococcaceae* (Fig. 4c), and other Firmicutes (Supplementary Fig. 5a), suggesting CWB2/SLH-loss and SH3-expansion represents their lineage-specific adaptation. Of the two species we studied experimentally, *Terrisporobacter glycolicus* and *Intestinibacter bartlettii,* only *I. bartlettii* exhibited evidence of S-layer presence (Fig. 2d, and Supplementary Fig. 4) and its SLLP contained the SH3 anchoring domain (Fig. 4a). Although the presence of SH3 domains is conserved across all *Terrisporobacter/Intestinibacter*, the SH3-anchored β-sandwich SLLP homologues were specific to a distinct group of *I. bartlettii* isolates (Fig. 2e) suggesting that this novel S-layer has evolved subsequently to the major changes to the cell envelope architecture in the *Terrisporobacter/Intestinibacter* ancestor.

For the anchoring domains to perform their function in the attachment of S-layers and other surface proteins to the cell, specific secondary cell wall polymers (SCWP) need to be synthesised and incorporated into the cell wall(*51*). In protein similarity network analyses of all proteins encoded in the 117 *Romboutsia*- and *Paeniclostridium*-genomes which have undergone multiple anchoring system swaps/gains/losses, we identified only a single protein without anchor enriched in CWB2-positive or SLH-positive species (Fig. 4d). The CWB2 system-associated protein was a homologue of a polysaccharide synthase (CD2777) involved in the synthesis of SCWP for CWB2 anchoring in *C. difficile*(*47*). The SLH system-associated protein was a homologue of TagA, involved in SLH-specific SCWP biosynthesis in Firmicutes, including *B. anthracis*(*52*) and *Paenibacillus alvei*(*51*). Therefore, enzymes required for the biosynthesis of SCWP anchors co-distribute with their cognate anchoring domains. The presence of these secondary cell wall polymer biosynthetic enzymes was also detected in organisms where both SLH and CWB2 proteins were encoded in the same genome (Fig. 4d). However, in those cases, the inferred SCWP type reflected that of the more abundant anchoring domain, while the second domain was found in only a few proteins (Fig. 4d), suggesting residual presence from incomplete anchoring domain swap, an indication that this can be a stepwise process. For S-layers to assemble on the cell surface, in addition to the presence of specific cell surface anchors, a protein secretion system is needed to transport SLLP monomers from the cytosol across the cell membrane. An SLLP-specific SecA2 secretion system is utilised in the assembly of α/β-sandwich *C. difficile* S-layer(*53*). However, our analyses revealed that neither of the *Peptostreptococcaceae* genomes encoding SH3- or CuOX-anchored β-sandwich SLLPs encoded any identifiable orthologues of SecA2. This suggests that, in *Peptostreptococcaceae*, SecA2 is associated predominantly with α/β-sandwich CWB2-anchored S- layers and some homologues of β-barrel SLH-anchored SLLPs, and that β-sandwich SH3- and CuOX-anchored SLLPs can utilise a more broadly distributed canonical SecA or a not yet identified alternative secretion system. Therefore, S-layers within a single bacterial family are not only assembled from different building blocks but also utilise different mechanisms for their assembly and the evolution of S-layer needs to be coordinated with associated cell wall modification and protein secretion machineries.

Together, our analyses revealed that S-layer anchor_DOM_s are part of broader cell wall anchoring systems shaped by modular evolution. Entire systems and anchor_DOM_ of individual proteins were swapped, acquired, or lost on multiple independent occasions, resulting in a mosaic of different anchors and functional domains often found in the genomes of closely related species. Homologues of enzymes known to be involved in the biosynthesis of CWB2- and SLH-specific secondary cell wall polymer anchors are commonly swapped alongside the SLH/CWB2 anchoring systems. This indicates that, at least in *Peptostreptococcaceae*, evolution of S-layer anchoring mechanisms is coordinated with that of cell wall chemistry. The capacity for utilisation of pre-existing swappable anchor_DOM_ modules can also provide an explanation for the diversity of anchors observed in S-layers sampled from across the Tree of Life, including those utilising the same surface_DOM_-fold (Fig. 1b).

### Extreme divergence, LGT, and recombination shape diversity of homologous S-layers

The observed diversity of S-layer cell anchoring mechanisms and variation in the number and type of individual subdomains within a single surface_DOM_ structural fold can be explained by modular evolution. However, at this point, it is unclear what mechanisms shape the structural and functional diversity of homologous S-layers utilising the same anchor_DOM_- and surface_DOM_-type. The extremely high level of sequence diversity of homologous S-layer surface-forming domains such as those of α/β-sandwich fold-type (Fig. 3b), complicates evolutionary analyses across the broader taxonomic scales. To circumvent this difficulty, and to better understand the mechanisms of the SLLP-surface_DOM_ evolution, we leveraged the established model of *C. difficile,* an important human pathogen where the S-layer provides a selective advantage within the host during infection(*30*).

Our bioinformatic analyses of available *C. difficile* SLLP (*Cd*SlpA) sequences (Fig. 5a) using Duplication-Transfer-Loss reconciliation identified signatures of at least 15 unambiguous recent LGT events (Fig. 5b), consistent with an active process, likely involved in immune evasion(*54, 55*). Of all 18,035 non-atypical *C. difficile* genome assemblies available in NCBI, 99.7% contained *Cd*SlpA coding genes. This suggests selection pressure for retention of a single copy of an *slpA* gene, in contrast to *Bacillus anthracis* with its two SLLPs(*6*), for example. Recombination analyses provided strong evidence for at least a single recombination event between parental SlpA_SLCT4_ and SlpA_SLCT6_, resulting in a recombinant divergent SlpA_SLCT4_ (Fig. 5c). Thus, recombination not only plays a part in exchanging different SLCT cassettes between different genome types but can also contribute to generating the diversity observed within a single species, such as among *Cd*SlpAs.

**Fig. 5:**
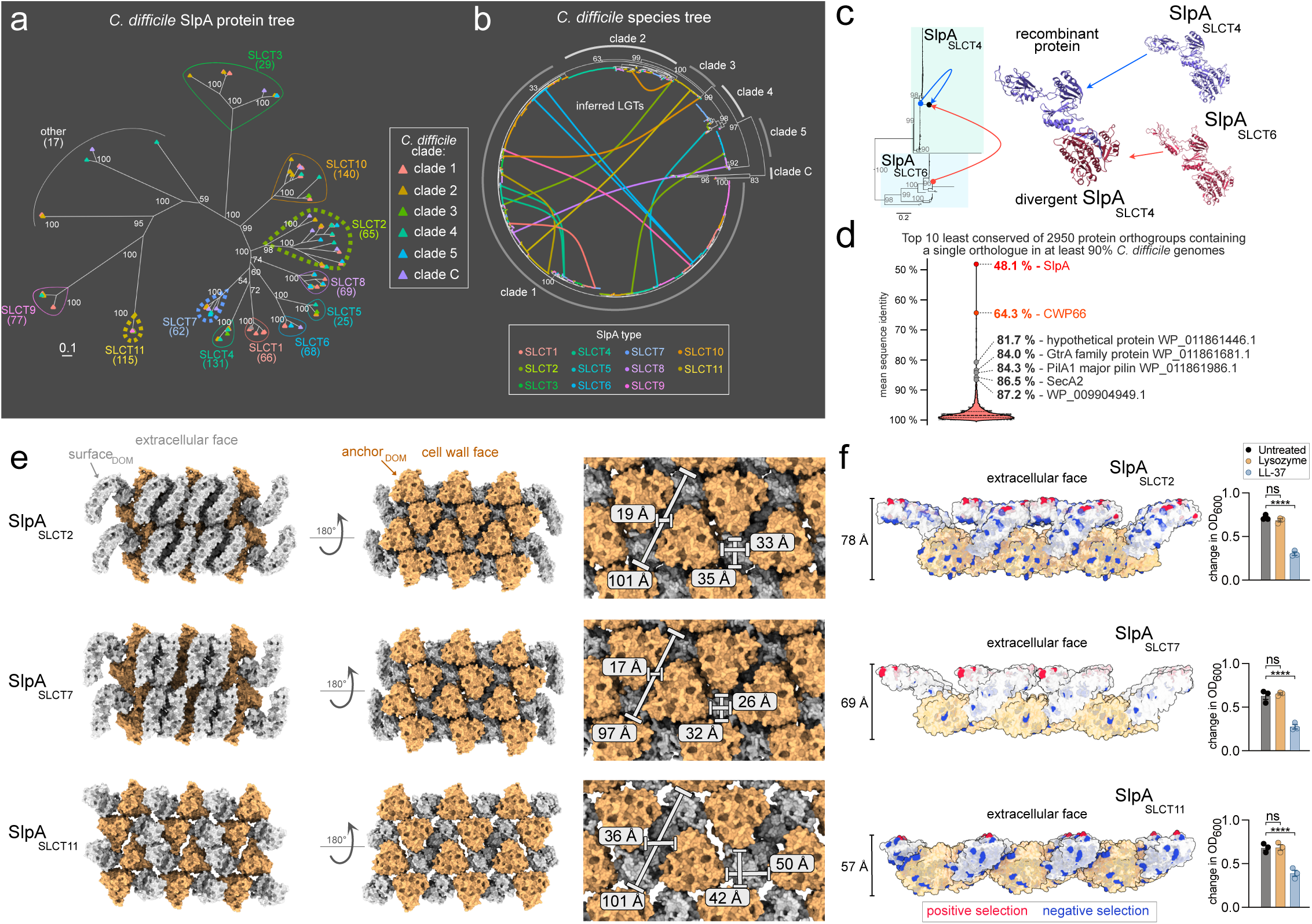
Evolutionary mechanisms shaping the diversity of S-layers across clinical isolates of a pathogen. **a,** Phylogeny of SlpA SLLP colour-coded based on the clade-membership of its source *C. difficile* genome assembly (tree tip colours). Diverse SlpA sequences belong to multiple distinct S-layer cassette types (SlpA_SLCT1-11_). **b,** The 14 unambiguous LGT events (coloured lines), identified using Duplication-Transfer-Loss reconciliation analysis, were mapped onto the core genome species tree of 866 *C. difficile* isolates including all six clades (clades 1-5 and C) and coloured based on the SlpA-type membership. **c,** An unambiguous recombination event (red and blue arrows) between SLCT4 and SLCT6 identified in recombination detection program RDP4 analyses was mapped onto a SlpA_SLCT4_ and SlpA_SLCT6_ protein phylogeny (left) and onto AlphaFold2 structure models (right) of the two inferred parent proteins (blue or red) and the recombinant sequence (red/blue). **d,** Violin plot represents the distribution of mean pairwise sequence identity within 2,950 orthogroups containing a single copy orthologue (SCO) in at least 90 % (240 of 267) analysed representative non-redundant *C. difficile* genome assemblies. Scale bars represent the number of substitutions per site. **e,** Representation of the SLLP-formed lattice derived from crystal packing of the three determined SlpA structures. The three SlpAs arrange in different lattice topologies, represented here at different tilt angles: extracellular (left) and cell wall-facing orientations (middle). Distances measured between neighbouring anchor_DOM_s (right, white lines) show that anchor_DOM_s are packed more closely in SlpA_SLCT2_ and SlpA_SLCT7_ lattices and are more separated in the SlpA_SLCT11_ lattice. **f,** Mapping of negatively (blue) and positively (red) selected residues onto a side view of the S-layer lattice shown in surface representation for each SlpA_SLCT_ (surface_DOM_ – white, anchor_DOM_ – orange, left). The three lattices differ in thickness reflecting the differences in structural organisation of the SlpA monomers. Despite their extreme sequence divergence (**d**) and structural differences (**e**), all analysed S-layers display conserved patterns of differentially distributed sites under adaptive evolution. While negatively selected residues are found in surface_DOM_ and anchor_DOM_ and on all SlpA surfaces, the sites under positive selection are exclusively found on the environment-exposed extracellular surface of the surface_DOM_. Broth-based killing assay (right) of different SlpA_SLCT_-carrying strains in the presence of lysozyme (orange) and LL-37 (blue) relative to the non-treated control (black) confirms conserved resistance to these innate immune effectors.

To investigate the level of SlpA divergence relative to other protein families encoded within *C. difficile* genomes, we identified 2,950 orthogroups containing a single copy orthologue in at least 90% (SCO90) of 265 non-redundant representative *C. difficile* genome assemblies. Within this dataset, SlpA was the least conserved of all identified SCO90s (48.1% average sequence identity, Fig. 5d). Thus, *C. difficile* SlpA represents the most rapidly evolving of all identifiable single-copy orthologue proteins found across all analysed *C. difficile* genomes. It is therefore likely that the high level of SlpA divergence reflects an adaptation of *C. difficile* to selective pressures inside the mammalian gut(*30*). This hypothesis is further supported by the observation that another major cell surface antigen, S-layer-associated protein Cwp66, previously implicated in adhesion(*56, 57*) and survival in the human gut(*57*), is the second least conserved SCO90 (64.3 % average sequence identity, Fig. 5d). The fourth least conserved SCO90 is a homologue of the cell wall teichoic acid glycosylation protein GtrA(*7*). *Listeria monocytogenes* homologue (GtcA) plays a role in cell wall glycosylation(*58, 59*), likely as a flippase(*60*), and the *B. anthracis* homologue is involved in the biosynthesis of SCWPs essential for S-layer surface attachment(*59*). The sixth SCO90 protein is SecA2, the secondary secretion system ATPase required for specific translocation of *Cd*SlpA(*53*), which co-distributes with CWB2-anchored SLLPs in *Peptostreptococcaceae* (Fig. 4c).

It is not clear if both LGT and high levels of sequence divergence can also contribute to the diversity of SLLPs in other bacterial lineages. Our bioinformatic analysis of *Bacillus* genomes revealed patterns consistent with both mechanisms in closely related *B. cereus/anthracis* isolates (Supplementary Fig. 7a-d), suggesting that these mechanisms are not limited to *C. difficile* SlpA and may be more common in other S-layer containing species. Interestingly, SAP/EA1 homologues were identified within the *B. cereus/anthracis*-group and not in their closest relatives from SAP/EA1-negative *B. cereus-*lineage (Supplementary Fig. 7d), suggesting differences in their cell envelope architectures. It is important to note that extreme levels of sequence divergence such as those observed in *Cd*SlpA (Fig. 5d) could mask more ancient LGTs, leading to underestimation of S-layer protein exchange between more distantly related taxa. In our analysis, we identified a single clear example of a β-sandwich EA1 surface-forming domain homologue being shared *via* LGT between *Bacillus* and *Peptostreptococcaceae* (Supplementary Fig. 7f,g). As *Peptostreptococcaceae* EA1 contained CWB2 and not SLH anchoring domains, it provided the first clear example of anchoring domain swap in SLLP homologues. We were able to experimentally verify the presence of an EA1 homologue in *C. mangenotii* LM2 surface protein preparations, where it constituted a minor S-layer component, alongside its dominant CWB2-α/β-sandwich SLLP (Supplementary Fig. 7h,i).

Together, our data demonstrate that the observed diversity of S-layers utilising the same anchor_DOM_- and surface_DOM_-types can be generated through extreme sequence divergence, homologous recombination, and LGT between close and more distant relatives. Our analyses also revealed patterns of accelerated evolution in S-layer-associated proteins and proteins involved in S-layer assembly.

### Extensive S-layer rearrangements and positive selection across clinical isolates of a pathogen

Over the broad phylogenetic scales, S-layer building blocks from the same structural family can assemble into diverse cell surface lattice architectures (Fig. 1b). To better understand the structural effects of the extreme SLLP sequence variation at the narrowest phylogenetic scales (Fig. 5d), we determined the crystal structure of *C. difficile* SlpA_SLCT11_ and SlpA_SLCT2_, representing distinct phylogenetic SlpA-types from multiple clinical isolates of a single species (Fig. 5a), and compared them to that of the previously studied SlpA_SLCT7_(*40*).

All tested SlpA homologues are proteolytically processed following secretion, resulting in two separate peptides: low molecular weight surface layer protein (LMW SLP or SLP_L_), containing the S-layer surface-forming domain, and high molecular weight (HMW SLP or SLP_H_), containing the CWB2 anchoring domain. Regardless of SLCT membership, the overall topology of *Cd*SlpA structures (Supplementary Fig. 8a) is maintained and consists of a triangular prism arrangement of the CWB2 anchoring domain, an interacting domain (ID) linking anchor_DOM_ and surface_DOM_, followed by one or two domains (D1-only, or D1/D2) of surface_DOM_, protruding the plane of the array(*40*). The structure of the anchor_DOM_ is highly conserved among all determined *Cd*SlpA structures, with core root mean square deviation (RMSD) of ∼1 Å (Supplementary Fig. 8a). This is consistent with the higher degree of anchor_DOM_ sequence conservation relative to that of surface_DOM_ in SLLPs from other organisms(*2*) and apparent in the analyses across the Tree of Life presented here (Supplementary Fig. 1-3). Within the more conserved *C. difficile* CWB2 anchor_DOM_, the highest degree of structure variability was observed in loops projecting from the anchor_DOM_ plane towards the extracellular face of the S-layer in SlpA_SLCT11_ or the surface_DOM_/anchor_DOM_ interspace in SlpA_SLCT2_ and SlpA_SLCT7_ (Supplementary Fig. 8a). Consistent with the more general pattern observed across the Bacterial Tree of Life (Supplementary Fig. 1 and 2), the most prominent differences across the SlpA structures were observed within their environment exposed S-layer surface-forming domain (Fig. 5e-f, Supplementary Fig. 8). Unlike the SlpA_SLCT7_ and SlpA_SLCT2_, which share a 2-domain (D1/D2) arrangement, SlpA_SLCT11_ lacks the entire D2 domain, which accounts for over 20% of total solvent accessible area of the S-layer unit, resulting in a smoother (Fig. 5e) and thinner array (Fig. 5f). The stark contrast between the surface_DOM_s of SlpA_SLCT2_, SlpA_SLCT7_, and SlpA_SLCT11_ provided evidence of how the entire fold that determines S-layer architecture can be reshaped within the relatively short phylogenetic timescales of emergence of different isolates of the same species. The *Cd*SlpA surface_DOM_ amino acid sequence regions align poorly with each other, further demonstrating their extreme sequence divergence (Fig. 3b and 5d) and providing a likely explanation for difficulties in identifying homologous SLLPs from more distantly related species. Interestingly, surface_DOM_-anchor_DOM_ interactions form the majority of the lattice interfaces in all studied *Cd*SlpAs (Supplementary Fig. 9), suggesting that the main role of the surface_DOM_ may be as a crystallisation enhancer module which forms an S-layer lattice only in conjunction with the more common CWB2 anchoring domain. This mode of S-layer lattice assembly, involving both the anchoring and surface domains, could explain the shorter surface_DOM_ in *Cd*SlpA relative to those identified in other *Peptostreptococcaceae* S-layer-types (Fig. 2c) and possibly the lack of any identifiable anchor_DOM_ swaps involving α/β-sandwich surface_DOM_.

Despite the observed differences in their lattice architecture, all the studied *C. difficile* isolates utilising different S-layer variants displayed conserved resistance to natural antimicrobial agents LL-37 and lysozyme produced by the human immune system (Fig. 5f). This demonstrates that considerable structural changes of lattice can evolve without compromising the physiological functions of the S-layer. It also raises important questions about the reasons behind the apparent sensitivity to the antimicrobials observed in other *Peptostreptococcaceae* utilising homologous S-layers (Fig 3d-f).

Finally, it remained unclear if the observed structural variation in *Cd*SlpAs is mainly shaped by genetic drift and recombination or if any of the sites are under positive selection. To test this, we analysed non-redundant nucleotide sequences encoding SlpA identified in all 18,035 analysed *C. difficile* genomes. All *Cd*SlpA types displayed patterns of positive selection and all identified sites under positive selection were within the surface_DOM_ and were surface exposed (Fig. 5f and Supplementary Fig. 10). This pattern was contrasted by the negatively selected residues found within surface_DOM_ and anchor_DOM_, and at all lattice-forming faces of the S-layer array (Fig. 5f). This is consistent with previous analyses indicating that surface antigens of pathogenic bacteria and viruses, and especially their exposed epitopes, are often under positive selection(*61, 62*).

The positive selection patterns detected only at the S-layer face exposed to the extracellular environment are consistent with its proposed role in phage and immune evasion. Extreme sequence and structural variation can facilitate the pathogen’s arms race with human adaptive immunity without compromising its conserved role in protection against lysozyme and LL-37, two components of the innate immune response. This model is consistent with the recently described capacity to generate monoclonal antibodies selective towards different SlpA variants(*63*).

In summary, the evolution of a single S-layer-forming protein can have a dramatic impact on the architecture of the bacterial cell surface at the narrowest phylogenetic scales which, in pathogens such as *C. difficile*, contributes to the protection against the human immune system.

## Discussion

As opposed to the archetypical elements of bacterial cell envelope architecture - lipid cell membrane and peptidoglycan cell wall - the main building block of a specific S-layer lattice is commonly encoded by a single gene and therefore can provide a substantial level of evolutionary plasticity to the microbial cell surface. This plasticity is of particular significance in bacterial pathogens such as *B. anthracis* or *C. difficile*, where the S-layer plays a role in disease(*10, 16, 30, 32*).

Our bioinformatic analyses demonstrate that the observed diversity of *C. difficile* S-layers is shaped by lateral gene transfer, homologous recombination, adaptive evolution, and extreme sequence divergence. Similar mechanisms can also generate the diversity of SLLP folds, and S-layer crystal lattice architectures across the bacterial Tree of Life (Fig. 1 and references(*34, 35*)), as we observed in wider sampling of *Peptostreptococcaceae* and *Bacillus*. Comparative analyses of the two crystal structures of *Cd*SlpA homologues experimentally determined in this study show that extreme surface_DOM_ sequence divergence can result in significant changes to the S-layer architecture including lattice thickness, anchor_DOM_ packing, and extensive rearrangements of its environment-exposed face. The observed architectural variation of *C. difficile* S-layers does not perturb its physiological functions in spore formation and resistance against effectors of the human innate immunity such as lysozyme and LL-37.

In the case of pathogenic bacteria, the extensive plasticity of their environment exposed surfaces may reflect an arms race with external threats to the pathogen’s survival such as bacteriophages(*18, 19*) and human adaptive immunity(*64*). Since the S-layer is an assembly of multiple copies of *Cd*SlpA, a single mutation of an environment-exposed positively selected residue can correspond to changes in up to an estimated 0.1 - 1% of the entire solvent accessible outer surface of *C. difficile* SlpA crystal lattice (Supplementary Fig. 9d). The observed structural plasticity, particularly at the more externally exposed domains (e.g. sites under positive selection), shows how a single mutation can affect a significant proportion of the host-pathogen interface. This includes altering potential surface epitopes, and therefore contribute to immune evasion, as *Cd*SlpA has been directly implicated in interactions with the host and eliciting immune response(*64*). Understanding the evolutionary mechanisms shaping *Cd*SlpA sequence diversity and the resulting structural plasticity of the S-layer lattice is important if we are to therapeutically target the S-layer(*10, 19, 33*). To avoid mutation and/or recombination-driven resistance mechanisms arising against any drugs designed to disrupt the S-layer, we need to consider the short-time scale evolution processes identified here. Furthermore, the structural landscape of S-layers needs to be considered as recent immunological studies identified that high-affinity anti-LMW monoclonal antibodies binding distinct epitopes of a single SlpA have different effects on *C. difficile*’s growth, toxin secretion, and biofilm formation(*33*).

The high level of S-layer diversity is not limited to clinical isolates of *C. difficile* and was observed in different strains of *Bacillus stearothermophilus*(*65*) and on a broader taxonomic scale in divergent SLLP sequences found across multiple species of *Bacillus* and *Lactobacillus*(*66*). At an even broader phylogenetic scale, among fourteen tested representatives of a single family of Bacteria, *Peptostreptococcaceae*, we detected and experimentally verified nine new SLLPs with one novel surface_DOM_ fold-type (β-barrel, CATH 2.4) and two anchoring domains (Cu-OX and SH3), not observed in any of the previously characterised SLLPs. Our results demonstrate that the diversity of S-layers in *Peptostreptococaceae* is shaped by modular evolution(*67, 68*). This is apparent when the mosaic of different genes, protein folds, secretion systems, and surface anchoring systems, consisting of different anchoring domains and different chemical pathways generating different secondary cell wall polymer modules, are mapped onto their species tree (Fig. 6a). However, this modularity is not restricted to a single mode of evolution but rather encompasses multiple ‘complexity’ levels. At the genome level, entire multigene S-layer cassettes can be swapped *via* LGT and recombination, and alternative multicomponent anchoring systems (e.g. CWB2 and SLH), including the surface-anchor biosynthesis machinery and its cognate anchoring domains of multiple proteins, can be replaced. At the individual protein level, alternative anchoring domain types can be swapped or gained, enabling otherwise homologous proteins to utilise one of the multiple cell-anchoring systems that coexist within a single bacterial genome (e.g. choline-binding and CWB2). Similar patterns were observed at the subdomain level in lattice-forming surface_DOM_s, as demonstrated by variation in the number of β-sandwich (sub)domains within Ig-like arrays(*69*), the most common fold of known SLLP architectures, acquisition of a novel α-helical surface subdomain in α/β-sandwich S-layer of *A. irregularis*, or as the capacity to lose the entire D2 subdomain in the α/β-sandwich fold of *C. difficile* SlpA_SLCT11_. In addition, the reorganisation of subdomains may also play roles in the incorporation of divergent S-layer proteins into an existing lattice such as that proposed for the minor S-layer component SlpX in *Lactobacillus*(*70*) or anchor-swapped EA1 homologue in *C. mangenotii.* At the cell biology level, the entire S-layer can be considered as a cell envelope module which can be swapped, remodelled, or lost. This multilevel modular architecture provides a flexible framework for various lineage- and species-specific cell surface adaptations (Fig. 6b). In *Romboutsia*, CWB2 and SLH secondary cell wall polymers were swapped, gained, or lost multiple independent times, often as part of major genome rearrangements leading to multiple inferred S-layer losses. In *Clostridioides/Asaccharospora*, a single core CWB2-anchored α/β-sandwich S-layer scaffold is retained, and the variation of the cell envelope is mostly due to the modification to the pre-existing lattice or addition of S-layer associated proteins. In *Terrisporobacter/Intestinibacter*, major ancestral changes to the cell wall chemistry have likely caused a loss or swap of the ancestral S-layer and driven the emergence of a novel SH3-anchored β-sandwich SLLP.

**Fig. 6:**
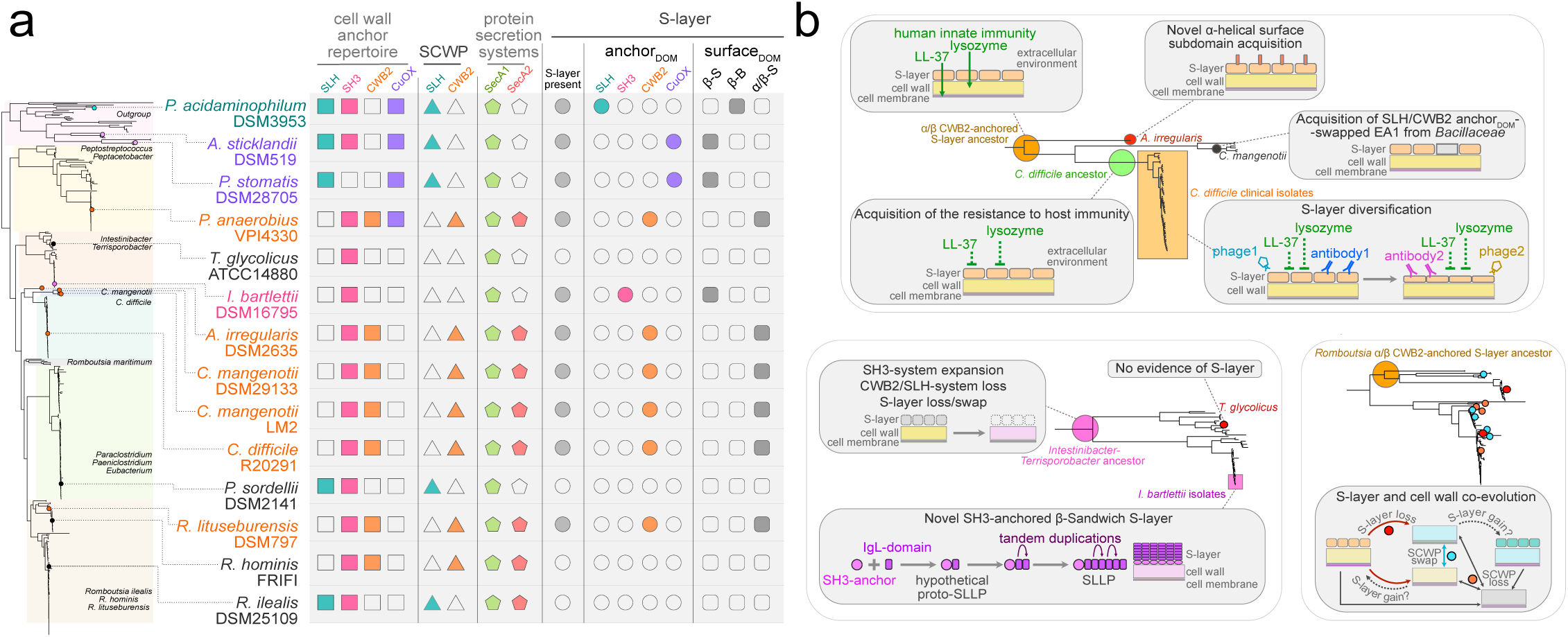
Multiple evolutionary pathways to S-layer exoskeleton gain, loss and diversification in a single family of Bacteria. **a,** An array representing presence (solid shapes) or absence (empty shapes) of cell wall anchoring domains found in at least a single protein per genome (rectangles), enzymes involved in biosynthesis of SLH- or CWB2-specific SCWP cell anchors (triangle), SecA protein secretion systems, including SecA2 involved in S-layer assembly in *C. difficile* (pentagons), experimentally verified presence of S-layer (grey circles), and anchor_DOM_-types (coloured circles) and surface_DOM_ folds (rounded rectangles) of experimentally identified SLLPs mapped onto a species tree of *Peptostreptococcaceae*. The scattered distribution of a mosaic of different cell envelope components including systems involved in S-layer assembly and anchoring to the cell is consistent with their diversity being shaped by modular evolution. **b,** Three distinct models of S-layer evolution based on the results presented in this study demonstrate that the observed diversity of S-layers is generated by multiple mechanisms underlying the extreme evolutionary plasticity of this major component of bacterial cell architecture: (i) structural and functional diversification of the conserved α/β-sandwich CWB2-anchored S-layers in *Clostridioides-Asaccharospora* lineage (top panel); (ii) recent strain-specific emergence of a novel SH3-anchored β-sandwich S-layer protein in *Intestinibacter bartlettii* (bottom left panel) and (iii) evidence of a specific loss of S-layer within *Romboutsia*, a lineage displaying a high frequency of cell envelope remodelling (bottom right panel).

Modular domain organisation was recently observed in the structure of the proposed S-layer-associated protein, extracellular Fe(III) reductase from the Gram-positive bacterium *Carboxydothermus ferrireducens*, containing a domain homologous to *B. anthracis* SLH(*71*). This suggests that modular evolution can play a role in additional functions provided by S-layer-associated proteins, in this case in the Fe(III) respiratory process, as well as in shaping S-layer architecture. Furthermore, modular architecture was observed in many cell wall binding proteins(*72*), including lytic enzymes containing choline-binding modules(*73*), consistent with our detection of cell wall homeostasis proteins being subject to frequent cell wall binding domain swaps. Therefore, the observed evolutionary mechanisms shaping the S-layer likely represent a more general phenomenon of bacterial cell envelope modularity that also includes cell wall and cell membranes. Although the gene clusters encoding proteins involved in peptidoglycan biosynthesis are highly conserved across Bacteria, rare insertions of outlier genes were observed in Firmicutes and Candidate Phyla Radiation (CPR)(*74*) and our results demonstrate that secondary cell wall polymer biosynthesis machinery was changed on multiple independent occasions, at least in Firmicutes. It was also hypothesized that another cell envelope module, the outer membrane, was lost in Gram-negative (diderm) bacteria multiple times, resulting in the independent emergence of Gram-positive (monoderm) lineages(*42, 75, 76*). However, as opposed to more complex cell membranes and cell walls, the main S-layer building block SLLP is encoded by a single gene and therefore it can provide an extreme level of evolutionary plasticity to microbial cell surfaces.

The diversification of S-layer cell surfaces can result not only in diverse architectures, symmetry types, and biochemical properties but also in variable physiological functions. Probably the most striking example is that of *Peptostreptococcaceae* S-layer utilising the same α/β-sandwich surface_DOM_-fold and the same CWB2 anchor_DOM_ where only human gut pathogens *P. anaerobius* and *C. difficile* isolates (with their diverse S-layer variants) are resistant to natural antimicrobials lysozyme and LL-37. In contrast, resistance to these innate immunity effectors was not observed in α/β-sandwich surface_DOM_-containing species unlikely to cause human disease including *A. irregularis* DSM2635 and *R. lituseburensis* DSM797 (isolated from mud), and *C. mangenotii* DSM29133 (isolated from intestine of a healthy mouse(*77*)). These results indicate that the presence of divergent versions of the same type of S-layer can contribute to the protection against the innate immunity effectors in pathogens but not in non-pathogenic species of the studied *Peptostreptococcaceae*. Therefore, S-layer evolution can play a role in adaptation to the pathogenic lifestyle. Furthermore, recent data suggests that distinct S-layers of *Lactobacillus* also play an important role in probiotic bacteria associated with optimal health, where it can act as a mediator of the host inflammatory response and prevent activation of pro-inflammatory signalling(*26, 28, 29*). It is currently unclear if these properties are conserved across diverse *Lactobacillus* S-layers(*66, 70*).

Our data also suggests that the adoption of certain S-layer architectures can enforce some structural and evolutionary constraints. While surface domains of some SLLPs appear to be found mostly with a certain type of anchoring domain, others have undergone multiple independent anchor_DOM_ swaps. For example, in our analyses, all the α/β-sandwich SLLPs, including *C. difficile* SlpA, contained only CWB2 anchoring domains. This could be due to structural or functional constraints such as the importance of the surface_DOM_-anchor_DOM_ interactions within their S-layer lattices, or it may merely represent a limitation of currently available sampling. One potential explanation for these observations is that the emergence of S-layers where surface_DOM_-anchor_DOM_ interactions form most of the lattice interfaces results in increased structural dependence between the specific surface_DOM_ and anchor_DOM_, leading to the entrenchment of this architecture. Such evolutionary ratchet mechanism(*78, 79*) could lead to a decreased capacity to accept alternative anchor_DOM_s. No such restrictions were observed in the more common β-sandwich surface_DOM_ homologues, which can function with multiple different anchor_DOM_s and can be transferred *via* LGT between more distantly related species.

Elucidating the origins and evolution of S-layers is complicated by the major gaps in our knowledge, including (1) insufficient number of experimentally verified S-layer-positive and -negative species from across the Tree of Life, (2) limited understanding of the conserved and lineage-specific S-layer functions outside of a few model organisms, (3) unclear capacity for the *de novo* origins of S-layers from non-S-layer proteins, (4) limited understanding of the physiological selection pressures and mechanisms underlying possible S-layer gains and losses. The limited experimental evidence for S-layer loss includes the emergence of *Cd*slpA mutants in response to the experimental exposure to specific antimicrobials (diffocins)(*10*), the absence of S-layer in some strains of *Corynebacterium glutamicum*(*36*), and interspersed phylogenetic distribution of S-layer positive *Peptostreptococcaceae* and their relatives with no evidence of S-layer presented in this study. Interestingly, pathogenic *Corynebacterium* species lack identifiable homologous of canonical S-layer protein PS2 found in the non-pathogenic representatives of the genus(*80*) and the presence of the S-layer in the pathogenic species awaits verification. If absent, this would suggest that, at least in some lineages, loss of the S-layer cell envelope module could also play a role in the emergence and evolution of pathogens. More indirect evidence of the possibility of S-layer loss comes from the absence of S-layers in model bacteria Gram-negative *Escherichia coli* and Gram-positive *Bacillus subtilis*(*2*). It is also unclear if any of the extant Eukaryotes contain cellular outer shell homologous to the prokaryotic S-layer.

Although the presence of S-layers *sensu stricto* to our knowledge has not been demonstrated in Eukaryotes, paracrystalline glycoprotein cell wall layers can be found in some species such as green algae *C. reinhardtii*(*81, 82*). The available evidence suggests that the S-layer is present in most *Archaea*(*2*), raising questions about whether it was also present in the Last Common Eukaryotic ancestor and subsequently lost or whether its loss was an important step in early Eukaryogenesis. Interestingly, the latest computational(*34*) and experimental(*83*) approaches did not identify S-layers and SLLP candidates in Asgardarchaeota, suggesting that the closest archaeal relatives of eukaryotes can be S-layer negative. However, this may not be a conserved feature of all Asgardarchaeota(*84*) and further experimental studies are required to robustly verify this hypothesis.

To our knowledge, there are no previously verified examples of S-layer gain by an S-layer-negative ancestor in the natural environment. However, further studies in *Peptostreptococcaceae,* including *Romboutsia,* or different species of *Corynebacterium*(*36, 80*) could provide a much-needed model for studying possible S-layer acquisitions. The presence of multiple bacterial species with extremely diverse S-layers and species with no evidence of S-layer scattered across *Peptostreptococcaceae* phylogeny presented in this study could lend credibility to the hypothesis that S-layers have evolved multiple independent times(*2*). Perhaps the most striking example is that of *I. bartlettii* where a lineage-specific SLLP has emerged within a subset of isolates of a single species, likely providing an excellent model for studying S-layer acquisitions. It remains to be tested if and how a non-S-layer protein can evolve *de novo* to form a paracrystalline cellular exoskeleton in nature. One captivating example of a possible evolutionary intermediate in the S-layer evolution is that of a toga, a proteinaceous surface layer with membrane patches, found in *Thermotoga maritima*(*85*). The evolutionary model of S-layer origin from an outer membrane or a toga-like cell envelope layer could be a reverse of that proposed for outer membrane biogenesis from the S-layer(*86*), i.e. a decrease in the proportion of lipids and an increase in the proportion of proteins capable of self-assembly within the outer cell envelope layer. This hypothesis, however, does not account for the possibility of the independent S-layer emergence in monoderm Bacteria, where, in the absence of the outer lipid-containing layer, *de novo* S-layer would have to evolve *via* a different mechanism. An alternative mechanism can be postulated based on the inferred origin of the S-layer in *I. bartlettii*. Its SLLP, containing six nearly identical β-sandwich Immunoglobulin-like surface_DOM_ subdomains, has emerged *via* multiple tandem duplications of a single ancestral subdomain of a type commonly found across bacteria. If this mechanism indeed plays a role in *de novo* evolution of S-layers, it could provide one explanation for the common use of β-sandwich fold in S-layers observed across the Tree of Life.

Our findings demonstrate that bacteria can rapidly gain, lose, replace, or modify the S-layer cell surface exoskeleton through the utilisation of a multilevel modular architecture. Thanks to this scalable framework, commonly composed of a single repeating SLLP building block, a single genetic change can significantly alter the entire exoskeletal lattice. The segmented design of the SLLP adds another layer of flexibility, allowing it to attach to various cell surface configurations through swappable cell anchoring domains. The ability to remove, add, or remodel surface domain segments promotes rapid adaptation of the S-layer layout. This versatile framework underlies the diversity of S-layer architectures observed across the prokaryotic Tree of Life and offers opportunities for bioprospecting new natural lattice designs for nanotechnology, protein engineering, and bio-product development. Our study uncovered the molecular, structural, and evolutionary processes behind multiple independent origins, losses, and the remarkable structural and functional divergence of this major component of bacterial cell biology. This extraordinary evolutionary plasticity underscores the frontline role of these biological exoskeletons in adapting to extracellular environments. In the case of pathogenic *Clostridioides difficile*, the S-layer has played a pivotal role in the emergence of ancestral resistance to innate immunity effectors and its evolvability contributes to the ongoing thug-of-war with adaptive immunity at the host-pathogen interface.

## Materials and Methods

### Inference of the Tree of Life

The Tree of Life was inferred as described previously(*87*), using the same dataset expanded with an additional 127 genomes of organisms with available evidence of S-layers and/or SLLPs, resulting in a final dataset of 2,705 Bacteria, 315 Archaea, and 149 Eukaryota genomes. Shortly, seven conserved single-copy marker proteins were identified using hmmsearch (-E 1e-5) of HMMER3.3(*88*). The identified orthologue sequences were aligned with MAFFT(*89*) trimmed with trimAL(*90*) (gappyout mode), concatenated with Biopython module Nexus(*91*), and used for the inference of phylogenies in IQ-TREE(*92*) under LG + G4 model with 1,000 ultrafast bootstraps(*93*).

### Peptostreptococcaceae, Bacillaceae and Firmicutes species trees

A set of 400 of all available NCBI(*94*) *Peptostreptococcaceae* genome assemblies, excluding overrepresented *C. difficile* genomes (18,035 assemblies analysed separately in this study), was downloaded and expanded by a minimal representative set of 53 *C. difficile* genomes, to include all SLCT-types and clades identified in our analyses, resulting in the final set of 453 *Peptostreptococcaceae* genome assemblies (Supplementary Data 2). To identify 111 broadly conserved *Peptostreptococcaceae* single-copy orthologues (Supplementary Data 3), protein sequences of 609 single-copy orthologues identified in our analyses of *C. difficile* genomes were used as queries in phmmer search (-E 1e-10) of HMMER3.3(*88*) against a local database of 1,264,675 protein sequences encoded in 453 analysed *Peptostreptococcaceae* genomes.

Due to a large number of available Firmicute genome assemblies (470,472), a custom set of 828 genomes (Supplementary Data 2), including representatives of all major lineages, was selected for further analyses. All 2,247,509 protein sequences encoded in the 828 Firmicutes genomes were analysed using EGN(*95*) protein similarity network analysis, and the largest network (1,442,176 proteins) was subdivided based on multilevel community detection algorithm in R. Of all identified protein networks, 84 represented single copy orthologues (Supplementary Data 3) broadly conserved across the analysed Firmicutes genomes.

In a dataset of all available 7,377 GeneBank^8^ *Bacillus* genome assemblies, the 84 Firmicutes single copy orthologues were identified (Supplementary Data 3) using phmmer search (-E 1e-10). As many *Bacillus* species are overrepresented, including multiple isolates of pathogens from *B. cereus/B. anthracis* lineage, a set of non-redundant assemblies was identified by pruning the redundant tree leaves (edge length = 0) from a species tree of 7,377 *Bacillus* using Treemmer(*96*) to 99.999999% of the original tree length (-RTL 0.99999999). The final set of 4,774 non-redundant *Bacillus* assemblies (Supplementary Data 2) encoding at least 75 of 84 orthologues was used for species tree inference.

For each dataset, protein sequences of the identified single copy orthologues were aligned with MAFFT(*89*) trimmed with trimAL(*90*) (gappyout mode), concatenated with Biopython module Nexus(*91*), and used for the inference of *Peptostreptococcaceae* (Supplementary Data 4), Firmicutes (Supplementary Data 5), and *Bacillus* (Supplementary Data 6) phylogenies in FastTree(*97*) under -lg -gamma model.

### Identification of the SLLP homologues and common domain sequence removal

To investigate the presence of known protein domains and sequence motifs in available experimentally verified SLLPs (Supplementary Data 1), the SLLP sequences were scanned against local copies of HMM profile databases SUPFAM(*98, 99*), Pfam(*100*), CATH(*101*), NCBI HMM(*102*) including TIGRFAM(*103*), and PROSITE(*104*) using hmmscan (-E 0.001) of HMMER3.3(*88*). To search for proteins and genomes containing the domains and sequence motifs identified in SLLPs, their HMM profiles were used in hmmsearch, HMMER3.3(*88*) against the Tree of Life dataset of 10,667,779 protein sequences encoded in the 3169 analysed genomes. To identify the SLLP homologues, their full-length sequences (Supplementary Data 1) and the surface_DOM_-only sequences (following anchor_DOM_ sequence removal, Supplementary Data 1) were used as queries in BlastP(*105*) searches (E-value threshold = 1e^-5^) against the Tree of Life dataset. Data presented in this paper indicate that cell surface anchoring domains can display broader phylogenetic distribution (Fig. 1a and Supplementary Fig. 1c and 2), be present in multiple otherwise unrelated proteins encoded within the same genome (Supplementary Fig. 3g), and share higher sequence and structural conservation than the domains responsible for the function of the anchored protein (e.g. surface_DOM_ of SLLPs, Fig. 3c and Supplementary Fig. 8a). This in turn can result in masking of their distinct evolutionary histories evident as clustering into a single protein similarity network of functionally and structurally unrelated functional domains sharing the same anchoring domain, and cause network ‘hairballs’ due to frequent anchoring domain swapping (Supplementary Fig. 3). To account for that, in addition to the full-length sequence and structural analyses, we analysed anchorless functional domain-only sequences separately (Supplementary Data 8). The sequence regions corresponding to the common anchor_DOM_s were identified with InterProScan 5(*106*) and removed from full-length protein sequences using Bio.Seq module of the Biopython package(*91*). Adjacent domain repeats (e.g. three subsequent CWB2 or SLH domains) were treated as a single region (the start and end of the removed region corresponded to the start of the first repeat and the end of the last repeat) and were removed from the full-length sequences including any in-between linker amino acid residues. Correct sequence removal was verified using InterProScan 5(*106*), manual sequence inspection, and AlphaFold2(*107*) structure predictions. For clarity, throughout the text, we collectively refer to the sequences left following the anchoring domain removal as the ‘functional domain’ sequence.

*Bacillus* SAP/EA1 SLLP homologues (Supplementary Data 7) were identified with the protein similarity network analyses of all *Bacillus* SLH/CWB2/SH3-containing proteins following the anchor_DOM_ sequence removal and verified with hmmsearch using custom HMM profiles generated with the identified SAP, EA1, and SAP2 surface_DOM_-only sequences.

### Protein similarity network analyses

Unless stated otherwise, all protein similarity network analyses were performed using the same protocol. Protein networks were generated with BLAST(*105*), and quick edge file optimisation implemented in EGN(*95*) using the following thresholds: BLAST (e = 1e^-5^), simple link parameter (E-value threshold = 1e^-5^, hit identity threshold = 20%, minimum hit coverage = 20%, without the best reciprocal hit condition enforcement), and with the quick edge file creation (E-value threshold = 1e^-5^, hit identity threshold = 20%, minimum hit coverage = 20%, minimal hit length = 30 amino acids). The identified networks were analysed using igraph(*108*) and dnet(*109*) packages in R(*110*) and visualised in Cytoscape(*111*). Where indicated, communities of the densely connected subgraphs were isolated using the multilevel and walktrap community detection algorithms implemented in the igraph package(*109*).

### Analysis of the cell anchoring domain-containing protein repertoires

To investigate the evolution of cell surface anchored protein repertoires, we used two separate datasets of organisms with evidence of frequent anchoring domain system swaps in our across the Tree of Life analyses: a broader taxonomic set of 828 Firmicutes genomes encoding 2,247,509 protein sequences, and narrower taxonomic set 453 *Peptostreptococcaceae* genomes encoding 1,264,675 protein sequences. Proteins containing CWB2 (PF04122.15), SLH (PF00395.23), and SH3 (PF08239.14) anchoring domains were identified using hmmsearch (-E 1e^-1^), HMMER3.3(*88*). To extract functional domain sequences (Supplementary Data 8), sequence regions corresponding to the anchoring domains (SLH, IPR001119; CWB2, IPR007253; SH3, IPR003646) were removed from the full-length sequences using Bio.Seq module of the Biopython package(*91*). The functional domain homology was investigated with EGN network analyses, annotated with the output of InterProScan 5, mapped onto species trees in R, and further verified with manual inspection of MAFFT(*89*) protein sequence alignments and structural prediction in AlphaFold2(*107*) for selected representatives.

To identify any additional homologues containing anchoring domains other than SLH, CWB2, or SH3 in *Peptostreptococcaceae*, including *Romboutsia*, anchor_DOM_-removed functional domain-only sequences from each functional domain homology network were aligned in MAFFT(*89*) and resulting alignments were used to generate custom functional domain HMM profiles. These profiles were used to search the original datasets with hmmsearch (-E 1e^-5^). The sequences identified with the functional domain HMM profiles (Supplementary Data 9 and 10) were analysed with EGN network analysis following anchor_DOM_ -removal (Supplementary Data 10), annotated with InterProScan 5^19^ to verify the presence of conserved functional domains and any additional anchor_DOM_s. The identified domains were mapped onto the networks in Cytoscape(*111*) and verified with manual inspection of MAFFT(*89*) protein sequence alignment and structural prediction in AlphaFold2(*107*) for selected representatives.

### Genomic region synteny

Initial genome alignments of *Romboutsia* and *Intestinibacter* genome assemblies downloaded from NCBI were performed with Mauve(*112*) to identify regions syntenic to those encoding SLLPs that we experimentally verified in S-layer-positive species. Final pairwise alignments for the identified genomic regions were generated using minimap2(*113*) with parameters (-X -N 50 -p 0.1 -c) and visualised in R using gggenomes(*114*) package.

### Identification of non-anchor_DOM_ proteins co-distributing with anchoring systems in *Romboutsia*

All protein sequences encoded in the available *Romboutsia* and *Paeniclostridium* genome assemblies (118), excluding those containing CWB2 (PF04122.15) and SLH (PF00395.23) anchoring domains identified using hmmsearch (-E 1e^-5^), were analysed using protein similarity network analysis. Analysed genomes were classified into two categories: CWB2-containing (23 assemblies encoding more than two CWB2 proteins) and SLH-containing (65 assemblies encoding more than two SLH proteins). Non-anchor_DOM_ proteins co-distributing with their respective anchor_DOM_ systems (Supplementary Data 10) were identified as protein networks containing sequences from more than 90% of the respective anchor_DOM_ -containing species (>21 CWB2 or >58 SLH) and no sequences from species containing the other anchor_DOM_ system.

### *Clostridioides difficile* genome assemblies used in bioinformatic analyses

Separate classification systems of *Clostridioides difficile* isolates are available(*37, 115*), including those based on core genome phylogenies of universally conserved gene sequences most likely inherited vertically, and classification based on the genomic cassettes shared between isolates *via* lateral gene transfer. To account for these distinct classifications representing different evolutionary processes shaping *C. difficile* isolate diversity, the following process was used to select representative genome samples. An initial set of *C. difficile* genome assemblies was selected based on previous analyses(*37, 115–119*) so that all major types, variants, and clades were included. This initial dataset was next expanded by including assemblies encoding divergent SlpA sequences which were not present in the original dataset and identified with BlastP and tBlastN(*105*) against the NCBI nr(*120*) and NCBI genome assemblies(*121*) databases, resulting in the 141 genomes dataset (Supplementary Data 2). Next, the 141 genomes dataset was further expanded by including a broad sample of genome assemblies from large-scale clinical isolate sequencing project(*122*) downloaded from EnteroBase(*123, 124*), selected based on examination of their clade- and SLCT-type phylogenies, resulting in the 866 genomes dataset (Supplementary Data 2). Within the 866 genomes dataset, no SlpA protein sequence was identified in a single assembly of *C. difficile* NT949(*125*), which in the core genome species tree grouped (100% bootstrap support) with SlpA-positive *C. difficile* CLO_CA2914AA_AS within clade1. However, tBlastN with SlpA from its closest relative revealed the presence of a frame-shifted SlpA sequence (NCBI locus tag BM535_18170) annotated as a pseudogene. This sequence is at the beginning of contig_808 (genebank MPDZ01000808.1), suggesting a possible incomplete sequence or assembly rather than SlpA sequence loss. The only genome assembly that encoded 2 SlpA sequences was *C. difficile* strain lsh09 (GCF_900688555.1), one of these sequences (SAMEA1402364_01087) - as in other *C. difficile* genomes - is located in a 515,328 bp long contig (CAADAX010000002.1) next to *secA2*, and the second partial sequence constituted of a short 412 bp (CAADAX010000091) contig, suggesting it may be a contaminant. Finally, we verified that our dataset was representative of known *C. difficile* diversity by analysing the complete set of 18,035 *C. diffcile* (Supplementary Data 2) genome assemblies deposited in NCBI GenBank, where we identified 18,238 gene sequences encoding SlpA homologues, including 845 non-redundant *slpA* gene sequences (Supplementary Data 7) belonging to the SlpA_SLCT_ groups (Supplementary Data 7) identified in our 866 genomes dataset. For genome assemblies without available predicted gene sequences, encoded genes were predicted using Prodigal(*126*). The SlpA homologues were identified using surface_DOM_-only query sequences of all SLLP types in tBlastN (-evalue 1e-5) searches. The dataset of all available NCBI *C. difficile* genome assemblies was downloaded from the NCBI GenBank database^8,40^ on 8th November 2023.

### Clostridioides difficile core genome species tree

To identify a set of one-to-one universal orthologues for *C. difficile* core genome species tree inference, an initial dataset of 866 genome assemblies sampled across all six clades and containing all SLCT representatives was analysed using sequence similarity network analyses, resulting in 4,089 networks.

Network decomposition following walktrap analyses resulted in a dataset of networks including 605 networks of one-to-one orthologues with average sequence identities ranging between 85.4 % and 100 %. Of these, the final set of 147 one-to-one orthologues (Supplementary Data 3) with sequence identity level below 98% was used for core genome species tree inference. The identified sequences were aligned with MAFFT(*89*), trimmed with trimAL (gappyout mode)(*90*) and resulting trimmed alignments were concatenated with module Nexus of the Biopython package(*91*) to a final alignment of 866 concatenated sequences of 37,580 amino acid sites, which was used to generate the *C. difficile* core genome species tree (Supplementary Data 11) with IQ-TREE(*92*) under LG + G4 model with 1,000 ultrafast bootstraps(*93*).

### Identification of Clostridioides difficile orthogroups

*Clostridioides difficile* orthogroups were identified using OrthoFinder(*127, 128*) with DIAMOND(*129*) sequence homology search option. To remove redundancy and reduce computational complexity for downstream analyses, the complete dataset of 866 genome assemblies was reduced to a minimal representative subsample of 267 assemblies based on careful examination of universally conserved orthologues and SlpA alignments and the core genome species tree and SlpA protein tree. OrthoFinder assigned 1,004,302 genes (99.8% of the total) to 7,762 orthogroups (Supplementary Data 12). Fifty percent of all genes were in orthogroups with 266 or more genes (G50 was 266) and were contained in the largest 1,782 orthogroups (O50 was 1,782). There were 862 orthogroups with sequences from all genome assemblies present and 609 of these consisted entirely of single-copy genes closely matching the 605 one-to-one orthologues identified in the full set of 866 genomes using protein similarity network analysis. To account for possible incomplete genome assemblies and sequencing errors such as those observed for slpA in *C. difficile* NT949 and strain lsh09 or rare cases of strain-specific gene loss and gain, we expanded the set of the 862 one-to-one orthologues to include orthogroups containing zero or more than one sequence in no more than 10% (27/267) genomes. This resulted in the final set of 2950 orthogroups containing a single copy orthologue in at least 90% (240/267) *C. difficile* genomes (SCO90, Supplementary Data 12). Mean sequence identities (Supplementary Data 12) were calculated as an average of trimAl(*90*) pairwise identity comparisons (-sident) on MAFFT(*89*) alignments of all sequences within each orthogroup.

### Lateral gene transfer detection

Lateral gene transfers were detected using the Duplication-Transfer-Loss reconciliation method implemented in Ranger-DTL 2.0(*130*). *Ranger-DTL* analysis was performed using an optimally rooted SlpA protein tree and a manually rooted species tree using default *Ranger-DTL* parameters in 100 replicates with different random seeds. The results from the replicates were aggregated using *AggregateRanger* with the default parameters. A single optimally rooted SlpA protein tree was identified using the *OptRoot* program with default parameters. The position of the *C. difficile* core genome species tree root was selected based on the published analyses(*131*), and our phylogenetic analyses of the available *Peptostreptococcaecae* genomes. A high level of sequence divergence in the Low Molecular Weight domain (LMW, which includes the surface_DOM_) results in high levels of uncertainty in its amino acid sequence alignments and low support values in inferred protein trees. To account for this uncertainty, we manually inspected species and protein trees and selected only unambiguous LGT candidates from distantly related species (from distinct species tree groupings), which grouped together in protein trees with high support.

### Detection of the amino acid sequence sites under adaptative evolution

Initially, amino acid sequences encoded by the 972 non-redundant nucleotide *slpA* genes identified across all 18,035 NCBI genomes, were aligned with MAFFT(*89*) and used to construct phylogeny with IQ-TREE(*92*) under best fitting (according to AIC and BIC)(*132*) WAG+F+I+G4 model with 1,000 ultrafast bootstraps(*93*). Due to the high level of sequence divergence, the accuracy of SlpA alignments within highly variable regions (mainly within LMW) was low. To account for that, we separately analysed SlpAs belonging to each SLCT category, where higher levels of sequence similarity enabled the generation of more reliable alignments. In addition, this enabled the use of each SLCT as a replicate in testing trends conserved across all SLCTs. SLCT types were extracted based on the SlpA protein tree topology. Amino acid sequences for each SLCT type were aligned separately in MAFFT(*89*), and were used to generate corresponding nucleotide alignment with a local version of PAL2NAL_v14(*133*), with the nogap option to trimm alignment regions with gaps. ReadAl(*90*) (-phylip3.2 option) was used to convert fasta files to phylip format. The resulting trimmed nucleotide alignments were translated to amino acid alignments with EMBOSS transeq 6.6.0.0(*134*). The trimmed amino acid alignments were used to reconstruct phylogenies for each SLCT with IQ-TREE(*92*) with the WAG+F+I+G4 model. Bootstrap values and branch lengths were removed for subsequent analyses in R using ape library(*135*). SLCT5 was excluded from the analysis as only five non-redundant sequences were available.

The SLCT nucleotide alignments and corresponding protein phylogenies were used in CODEML analyses of PAML package(*136*) under M0 (one-ratio), and M1a (NearlyNeutral) and M2a (PositiveSelection) site models with heterogenous ω across the sites using following parameters: runmode = 0 (unrooted user tree), seqtype = 1 (codons), CodonFreq = 7 (FmutSel substitution model(*137*)), clock = 0 (no molecular clock assumption), model = 0 (sites models with homogenous ω across the branches). The different models were compared with the likelihood ratio test as described in the published protocol(*138*). Positively selected residues were identified in Bayes Empirical Bayes (BEB) analysis(*139*) with probability P > 95%.

Recombination can complicate the interpretation of the dN/dS analyses, and although we identified only a single unambiguous example of recombination within our dataset, we used GARD to further account for possible recombination events(*140*). The relative simplicity of the GARD approach had the added benefit of accounting for the apparent heterogeneity in modes of evolution observed between the cell wall anchoring domains (e.g. CWB2) and the functional domains (e.g. surface_DOM_) i.e. often the inferred breakpoints corresponded to the split between the two domains. Output files of the SLCT nucleotide alignment processed with GARD were used in all subsequent analyses (BUSTED, FUBAR, and MEME). FUBAR(*141*) was used to infer the positively (posterior probabilities > 90%) and negatively (posterior probabilities > 98.5%) selected sites. More inclusive thresholds were used for positive selection as it was verified with two additional methods (MEME and PAML), while more stringent thresholds were used for the negative selection identified using only a single method. BUSTED(*142*) was used to test for gene-wide episodic positive selection in at least one site on at least one branch while accounting for across the sites synonymous substitution rate variation(*143*). MEME was used to identify evidence of specific sites undergoing episodic diversifying selection (p-value < 0.1)(*144*). Finally, sites identified using all three methods (PAML, FUBAR, MEME) were mapped onto our experimentally determined *Cd*SlpA crystal structures and AlphaFold2(*145*) models of the remaining *Cd*SlpA homologues. Only sites identified with at least two of the three methods (PAML, FUBAR, MEME) were considered to display strong evidence of positive selection (Supplementary Data 13).

### Detection of recombination between *Clostridioides difficile* SlpAs

Evidence of recombination across the SlpAs sampled from all available *C. difficille* genomes was detected using RDP4.101 and RDP5 Beta 5.53(*146*), with no differences in the results obtained with each version. The 972 non-redundant nucleotide *slpA* genes identified across all 18,035 NCBI genomes and their corresponding MAFFT(*89*) amino acid alignment were used to generate nucleotide alignment with a local version of PAL2NAL_v14(*133*). The resulting nucleotide alignment was used as an input in a full exploratory recombination scan using RDP(*147*), GENECONV(*148*), Bootscan(*149*), Maxchi(*150*), Chimaera(*151*), SiSscan(*152*), and 3Seq(*153*) methods with default parameters. Of all 40 candidate recombination events detected using at least one of the seven methods, 9 candidate events were statistically significant in all seven methods, of which both inferred parental sequences were ‘known’ in four events (i.e. their sequences were present in the tested dataset). Of the four strongly supported recombination events with both parent sequences available, only a single unambiguous recombination event was identified following the manual verification within SlpA protein phylogeny and nucleotide sequence alignments of the parent and recombinant sequences. Recombinant sequence: GenBank=CP068558.1, locus_tag=JMW52_06215, protein_id=QQY54376.1; Major parent: GenBank=PRZC01000013.1, locus_tag=C4007_03100, protein_id=MDB2714594.1;

Minor parent: GenBank=ABLTPH020000028.1, locus_tag=REJ96_016885, protein_id=MDS6367270.1. Significance values from RDP4 analysis: RDP (2.6 E-58), GENECONV (5.9 E-54), Bootscan (1.5 E-60), Maxchi (4.3 E-24), Chimaera (4.1 E-23), SiSscan (4.9 E-35), 3Seq (5.6 E-11).

### Other bioinformatics methods

Unless stated otherwise, protein phylogenies were inferred using sequences aligned with MAFFT(*89*) in IQ-TREE(*92*) with 1,000 ultrafast bootstraps(*93*) under best fitting model selected with ModelFinder(*132*) according to Bayesian information criterion implemented in IQ-TREE. Multiple sequence alignments were inspected using Jalview(*154*). Phylogenetic trees were inspected in Archaeopteryx(*155*). Other general bioinformatics packages used for data analyses and visualisation included: ggplot2(*156*), dplyr(*157*), ggmsa(*158*), ggtree(*159*), ape(*160*), treeio(*161*), ggtreeExtra(*162*), ggnewscale(*163*).

### General microbiology methods

*Peptostreptococaccceae* strains were selected based on phylogenetic analyses and obtained from reference culture collection (DSMZ, German Collection of Microorganisms and Cell Cultures GmbH) or individual laboratory sources. All bacterial strains and specific culture conditions used in this study are described in Supplementary Table 2. Bacteria were cultured in DG250 or A35 workstations (Don Whitley Scientific) in anaerobic conditions (10% H_2_, 10% CO_2_, 80% N_2_).

### Bacterial growth

To investigate the presence of S-layer and expression of S-layer proteins in different culture conditions, all strains were cultured in strain-specific media following DSMZ strain collection protocols (Supplementary Table 2) and three media commonly used for the model *Peptostreptococcacea* bacterium *C. difficile*: TY (3% tryptose, 2% yeast extract), BHIS (brain heart infusion, 0.5% yeast extract, 0.1% L-cysteine), and chemically defined minimal medium (CDMM(*164*)). Bacterial growth was tested at 34 °C and 37 °C for all media and strains, and additionally at 28 °C for *C. mangenotii* LM2. All cultures were started by inoculating fresh media with overnight pre-cultures, in Hungate-type tubes to an OD_600nm_ of 0.08 measured in WPA CO8000 Cell Density Meter. To test the presence of S-layer and SLLPs at different growth phases, samples were taken at early-, middle-, and late-exponential, and early-, and late-stationary phases time points and OD_600nm_ values indicated in the results. Supplementation of growth media with CaCl_2_ to a final concentration of 500 μM was carried out to assess the impact of calcium on S-layer proteins in culture.

### CRISPR interference for targeted knockdown of *sllp* gene expression

*E. coli*-*C. difficile* shuttle vector for CRISPR interference, pIA33(*165*), kindly provided by Prof. Craig Ellermeier, was used to target expression of genes encoding identified SLLP in *R. lituseburensis* and *C. difficile*. The sgRNA was analysed using CASPER(*166*) (https://github.com/TrinhLab/CASPERapp), generated by amplification using oligonucleotide pairs: pIA33_ SLLP_2_Rlitu_F and pIA33_Insert_R; pIA33_SlpA1_F and pIA33_Insert_R listed in Supplementary Table 3 using pIA33 template, to substitute sgRNA-rfp, and introduced into MscI+PstI-digested pIA33 backbone by isothermal assembly(*167*). Negative-control plasmid (pIA34(*165*)) containing non-targeting sgRNA sequence that does not recognize any sequence in the genome, was used to control for non-specific plasmid effect. CRISPRi::*sll*p plasmids were introduced into *Peptostreptococacceae* host by conjugation(*168*). Transformants were selected on TY thiamphenicol 10 µg mL^-1^ agar, supplemented with cycloserine (250 µg mL^-1^).

Induction of dCas9-opt activity was mediated by supplementation of growth media with xylose (1% w/v) and the interference was observed for *C. difficile slpA* (as shown before(*165*)) and for *R. lituseburensis sllp*, as demonstrated by SDS-PAGE of surface layer extracts, RT-qPCR, and electron microscopy.

Viability of CRISPRi::*sllp* knockdown strains was assessed by spot assay, where a 5 µl aliquots of serial 10-fold dilutions of xylose induced and non-induced overnight cultures, were spotted onto TY-agar. Following 24-hour incubation, CFU mL^-1^ were enumerated.

A RT-qPCR was performed to confirm gene silencing, using oligonucleotide pairs: qCdR20291_SlpA_F and qCdR20291_SlpA_R; qCdR20291_RpoC_F and qCdR20291_RpoC_R; qCdR20291_GyrA_F and qCdR20291_GyrA_R; qRlitu_SLLP_F and qRlitu_SLLP_R; qRlitu_RpoC_F and qRlitu_RpoC_R; qRlitu_GyrA_F and qRlitu_GyrA_R, designed with PrimerQuest (Integrated DNA Technologies) listed in Supplementary Table 3. RNA extraction was carried out with Monarch Total RNA Miniprep kit (NEB) from 8 h cultures. DNAse treatment was carried out with TURBO DNA-free™ Kit (Invitrogen). cDNA preparation was performed using the SuperScript III VILO cDNA synthesis kit. RT-qPCR was performed using Luna Universal qPCR master Mix (NEB) in LightCycler96 (Roche). Samples of two biological replicates were analysed in 3 technical replicates. Results were calculated by the comparative cycle threshold method(*169*) and normalized to the gyrA transcript. Relative expression of *sllp* and *rpoC* control was presented as a log2 fold change, relative to non-induced CRISPRi::*sllp* expression level.

### Broth-based killing assay

Human innate immune effector (lysozyme; 500 µg mL^-1^, or LL-37; 5 µg mL^-1^) was added to mid-exponential cultures (OD_600nm_ ∼0.5) and its impact on the growth of strains expressing intact SLLP or of CRISPRi *sllp* knock-down was assessed by monitoring OD_600nm_. Two biological and three technical replicates were analysed in microplate reader (Stratus, Cerillo). OD change was estimated from a difference between OD_600nm_ at endpoint (8 h timepoint for *C. difficile* and 5 h for other *Peptostreptococacceae*) and OD_600nm_ measured upon supplementation with the additive.

### Sporulation assay

CRISPRi::*sllp* knock-down strains of *C. difficile* and *R. lituseburensis* presented sporulation defects compared to negative control when assessed by electron microscopy of whole cells. To estimate sporulation efficiency, the knock-down strains were cultured overnight, in biological duplicate. Aliquots of 50 µl were spotted onto selective TY-agar supplemented or non-supplemented with xylose and cultured for 24 h. Growth was collected from the plate and resuspended in 500 µl PBS. Cell suspension was split and one aliquot of 250 µl was heat-treated at 70 °C for 20 min. Both non-treated and heat-treated aliquots were serially 10-fold diluted in PBS, and 5 µl aliquots of dilutions were spotted onto TY, 0.1% taurocholate, 0.1% cysteine agar plates. Following overnight incubation, CFU mL^-1^ was enumerated. The heat-resistant CFUs are represented as a percentage of total viable (non-treated) CFUs.

### Transmission electron microscopy (TEM) of bacterial cell sections

Stationary phase bacterial cultures were harvested by centrifugation at 2,000 x*g* for 10 min at room temperature. Cell pellets were collected in 1 ml of fixative (2% glutaraldehyde in 0.1 M sodium cacodylate buffer, pH 7.4) and stored at 4 °C until further processing, performed at Newcastle University Electron Microscopy Research Services (EMRS). Samples were enrobed in low melting point agarose and post-fixed in 1% osmium tetroxide, dehydrated in gradient acetone, and embedded in epoxy resin.

Cell surface glycopolymers were visualised using a published protocol(*47*). Briefly, bacterial colonies were collected from 24 h agar plate cultures and pre-fixed with 2% glutaraldehyde and 10 mM L-lysine in CB-RR (0.1 M sodium cacodylate buffer, pH 7.4, 0.1% ruthenium red) for 10 min at room temperature.

Subsequently, pellets were fixed in 2% glutaraldehyde in CB-RR for 2 h at room temperature, washed with CB-RR, and post-fixed in 1% osmium tetroxide in CB-RR. Samples were then enrobed in agarose, dehydrated in a gradient of ethanol, and embedded in the polyhydroxy aromatic acrylic resin LR white.

Ultrathin sections of around 70 nm were deposited on copper grids and stained with uranyl acetate and lead citrate. Grids were then visualised using a Hitachi HT7800 120 kV transmission electron microscope (EMRS, Newcastle University).

Line profile analysis of cell envelope across cell sections was performed with Fiji(*170*).

### Electron microscopy of isolated S-layer lattice fragments

S-layer preparation for electron microscopy was performed as previously described(*40*) with minor modifications. Mid-exponential cultures were harvested by centrifugation at 2,000 x*g* for 15 min at room temperature. Cell pellets were washed twice in ice-cold deionised water (0.25x of culture volume) and disrupted by pulse-sonication for 2 min on ice. Unbroken cells were removed by centrifugation at 800 x*g* for 10 minutes at 4 °C and the supernatant was further centrifuged at 3,000 x*g* for 10 minutes. Harvested pellets containing S-layer fragments were washed with ice-cold 1 M NaCl (to an effective OD_600nm_ of 100, of initial culture) and resuspended in cold 2% Triton X-100 (to an effective OD_600nm_ of 200, of initial culture).

Preparations were loaded on glow-discharged, carbon-coated 300 mesh copper EM grids (Agar Scientific), stained with 2% uranyl acetate, and assessed using a Hitachi HT7800 transmission electron microscope operating at 120 kV (EMRS, Newcastle University).

For electron cryo-microscopy, S-layer fragments were diluted 10-fold in cold deionised water and added onto plasma-treated Quantifoil Cu R2/2, 200 mesh grids. The grids were blotted for 5 seconds at 4 °C, 90% relative humidity, vitrified in liquid ethane, using Vitrobot (FEI), and stored in liquid nitrogen.

Cryo-EM grid preparation and imaging were performed at the York Structural Biology Laboratory (YSBL) on a Glacios electron microscope equipped with a Falcon 4 direct electron detector (Thermo Fisher Scientific). Micrographs were taken at 200 kW with a magnification of 120,000 and −2μm defocus. Images were viewed and Fourier transforms were generated using 3dMOD(*171*) and Fiji(*170*).

### Identification of S-layer lattice proteins

As the main building blocks of known bacterial S-layers, SLLPs typically constitute one of the most abundant cellular polypeptides and the dominant protein in S-layers, aiding their identification from dominant bands in gel electrophoresis of S-layer preparations.

To identify the *Peptostreptococcaceae* SLLPs, bacteria with evidence of S-layer presence in electron micrographs of S-layer fragments and whole cell sections were cultured to stationary phase and harvested by centrifugation at 4,000 x*g* for 10 min at room temperature. Additional experiments with sampling from across the growth cycle were performed for *C. mangenotii* LM2, which contains both CWB2-SLLP and CWB2-EA1, and for the strains where no evidence of S-layer was found in the electron micrographs.

Harvested wet cell pellets were washed with a HEPES-NaCl buffer (50 mM HEPES, 100 mM NaCl, pH 7.5), followed by a critical step of the careful removal of all HEPES-NaCl liquid to ensure consistent extraction conditions in the subsequent steps. Non-covalently bound proteins, including S-layer proteins, were extracted from the cell surface using three different chemical methods. Preparations using 0.2 M glycine pH 2.2 were carried out as previously described(*40*). Two additional methods of extraction: 6 M urea, 50 mM Tris pH 8.0, 1 mM phenyl-methyl-sulphonide-fluoride (PMSF); and 1% sodium dodecyl sulphate (SDS), 50 mM Tris–HCl pH 8.0, 1 mM PMSF were used following the published protocols(*172, 173*). For each method, the washed cell pellets were resuspended in extraction solution to an effective OD_600nm_ of 50 and incubated at room temperature for 30 min, followed by centrifugation at 21,000 x*g* for 10 min. A 2 M Tris base was used to neutralise the pH of low-pH glycine preparations. Extracts were then mixed with a 0.25x volume of sample buffer (8% SDS, 400 mM DTT, 40% glycerol, 200 mM Tris-HCl pH 6.8, 0.4% bromophenol blue) and separated via 6-20% gradient SDS-polyacrylamide gel electrophoresis (SDS-PAGE).

Recovered spent media were used to assess secreted protein content and cell autolysis, by precipitation with 10% (v/v) of trichloroacetic acid (TCA), for 30 min on ice. Following centrifugation at 21,000 x*g* for 10 min at 4 °C, the precipitate was washed 3 times with ice-cold 90% acetone. The resulting air-dried pellet was resuspended in PBS to an effective OD_600nm_ of 50, mixed with a 0.25x volume of sample buffer (8% SDS, 400 mM dithiothreitol (DTT), 40% glycerol, 200 mM Tris-HCl pH 6.8, 0.4% bromophenol blue) and separated via 6-20% gradient SDS-PAGE.

To identify proteins in S-layer preparations, trypsin digests of the respective bands excised from SDS-polyacrylamide gels were analysed by LC-MS/MS in positive ion mode on a Finnigan LTQ ion trap mass spectrometer (Thermo Fisher Scientific, Bremen, Germany). Protein identification was performed using the Mascot MS/MS search engine program (Matrix Science Ltd, London) against a custom protein sequence database (*Peptostreptococcaceae*, containing 179,609 unique sequences created on 26/01/2023 from UniProt).

### X-ray crystallography of SlpA

TY broth cultures (16 h) of *C. difficile* strains were centrifuged at room temperature at 5,000 x*g* and the resulting pellets were washed with 0.1x culture volume of 50 mM M HEPES pH 7.4, 150 mM NaCl buffer. S-layer extraction was performed by resuspending the washed pellet in 0.01x culture volume of 0.2 M glycine-HCl pH 2.2, as described above. S-layer extract was filtered and resolved onto a Superdex 200 26/600 column using an ÄKTA Pure FPLC system (Cytiva) in 50 mM Tris-HCl pH 7.5, 150 mM NaCl buffer. Purified proteins were concentrated to 10 mg mL^-1^ and subjected to crystallization with the sitting drop vapour-diffusion method at 20 °C with a range of commercial screens.

Although SlpA_SLCT11_ was purified from Ox247 (wild type strain(*37*)) and Ox247ι1orf2 (mutant defective in glycosylation of SlpA(*174*), denoted SLCT11*), only the non-glycosylated protein resulted in successful crystallisation in 4 mM oxometalates, 50 mM Tris (base); 50 mM bicine 8.5, 20% v/v glycerol; 10% w/v PEG 4000 (Morpheus Fusion, Molecular Dimensions). SlpA_SLCT2_ purified from Opt2472 strain(*122*) produced diffraction quality crystals in 1.6 M Sodium citrate tribasic (Wizard Classic 3, Molecular Dimensions).

Diffraction data were collected on I24 beamline (λ = 0.71 Å) at the Diamond Light Source Synchrotron (Didcot, UK; mx24948-136) at 100 K. The data was processed with xia2(*175*) (SlpA_SLCT11*_) or DIALS(*176*) (SlpA_SLCT2_), scaled with Aimless(*177*) within CCP4 Cloud(*178*). Initial phases were obtained by molecular replacement in Phaser(*179*) with models of the CWB2 anchoring domain using the SlpA_CD630_ (PDB ID: 7ACY) template for SlpA_SLCT11*_ or the AlphaFold2(*145*) predicted model for SlpA_SLCT2_. The generated solution model was then subjected to automatic model building with ModelCraft(*180*), followed by iterative cycles of manual building with Coot(*181*) and refinement in Refmac5(*182*).

Final models were obtained after iterative cycles of manual model building with Coot and refinement in Phenix_refine(*183*) and PDBREDO(*184*). Validation of final models was performed using Coot and Phenix internal tools, as well as MolProbity(*185*) web server. Data collection and refinement statistics are summarized in Supplementary Table 4.

### Other structural analyses

Structural prediction for proteins of interest was performed with ColabFold v1.5.5: AlphaFold2 using MMseqs2(*107*). Comparative structural searches were carried out with Foldseek(*186*) and Dali(*187*) servers. Structural analyses including crystal contacts, distances, angles and dimensions were performed with PDBSum(*188*), PISA(*189*) and with plugins implemented in PyMOL Molecular Graphics System (Schrödinger, LLC) or UCSF ChimeraX(*190*). Graphical representations of S-layer proteins and crystal latices were generated using UCSF ChimeraX. SymProFold pipeline was used to predict S-layer lattice assemblies(*35*) following the standard protocol (https://github.com/symprofold/SymProFold_Tutorial_Data). AlphaFold3 server, using the PDB2025 (up to 3 February 2025) template option, was used to generate structural predictions for full-length and surfaceDOMs models for experimentally identified SLLPs across *Peptotreptococcaceae.* Individual subdomains – anchor_DOM_, surface_DOM_, D1, D2 and ID - were manually extracted from the ful-length predictions and aligned using the LSQ function in COOT. Calculated RMSD for pairwise superimpositions in Coot were represented as a heatmap using Rstudio.

## Acknowledgements

This work was supported by a Medical Research Council grant [MR/V032151/1] awarded to P.S.S and A.B.-S. and a Research Excellence Development Award [NU-018171] from the Faculty of Medical Sciences, Newcastle University awarded to A.B.-S. We thank Dr Joseph Gray for expert mass spectrometry analysis and protein identification, and Prof Craig Ellermeir for providing plasmids used in CRISPRi. Our thanks to Tracey Davey and Ross Laws from Newcastle University Electron Microscopy Research Services, supported by BB/R013942/1 for TEM sample processing; York Structural Biology Laboratory, University of York, supported by the Wellcome Trust (206161/Z/17/Z) for supporting cryo-EM data collection; Newcastle University Structural Biology Laboratory for facilitating crystallisation and data collection. We thank Diamond Light Source for access to beamline I24 (mx24948-136), through the “Macromolecular Crystallography at Newcastle, Durham, Lincoln and Durham” BAG). The contents of this work are solely the responsibilities of the authors and do not reflect the official views of any of the funders, who had no role in study design, data collection, analysis, decision to publish, or preparation of the manuscript.

## Author contributions

K.M.S., A.B.-S. and P.S.S. are the corresponding authors. K.M.S. and A.B.-S conceptualised and planned the study. K.M.S. wrote the manuscript with contributions from A.B.-S. and P.S.S. K.M.S. performed all bioinformatic analyses. A.B.-S and K.M.S. performed experimental work. A.B.-S crystallized *Cd*SlpA. A.B.-S. and P.S.S. determined and analysed protein structures. P.S.S. and A.B.-S. secured funding. K.M.S. and A.B.-S. managed the project.

## Competing financial interests

The authors declare no competing interests.

## Data availability

The authors declare that the main data supporting the findings of this study are available within the article and Supplementary information. Data sets generated and analysed in this study are provided as Supplementary Data Files. The atomic coordinate files for X-ray structures have been deposited in the Protein Data Bank (PDB) with the accession codes 9F8E, 9F8F.

## Notes

### Competing Interest Statement

The authors have declared no competing interest.

### Summary of Updates

Figures in the previous version had poor resolution after PDF conversion so an updated file with higher resolution is now being submitted.

