## Supplementary Information for "Evolutionary plasticity of bacterial surface layer protein exoskeletons"

File contains:
Supplementary Figures 1-10
Supplementary Tables 1-4
Supplementary references
Supplementary Data Inventory

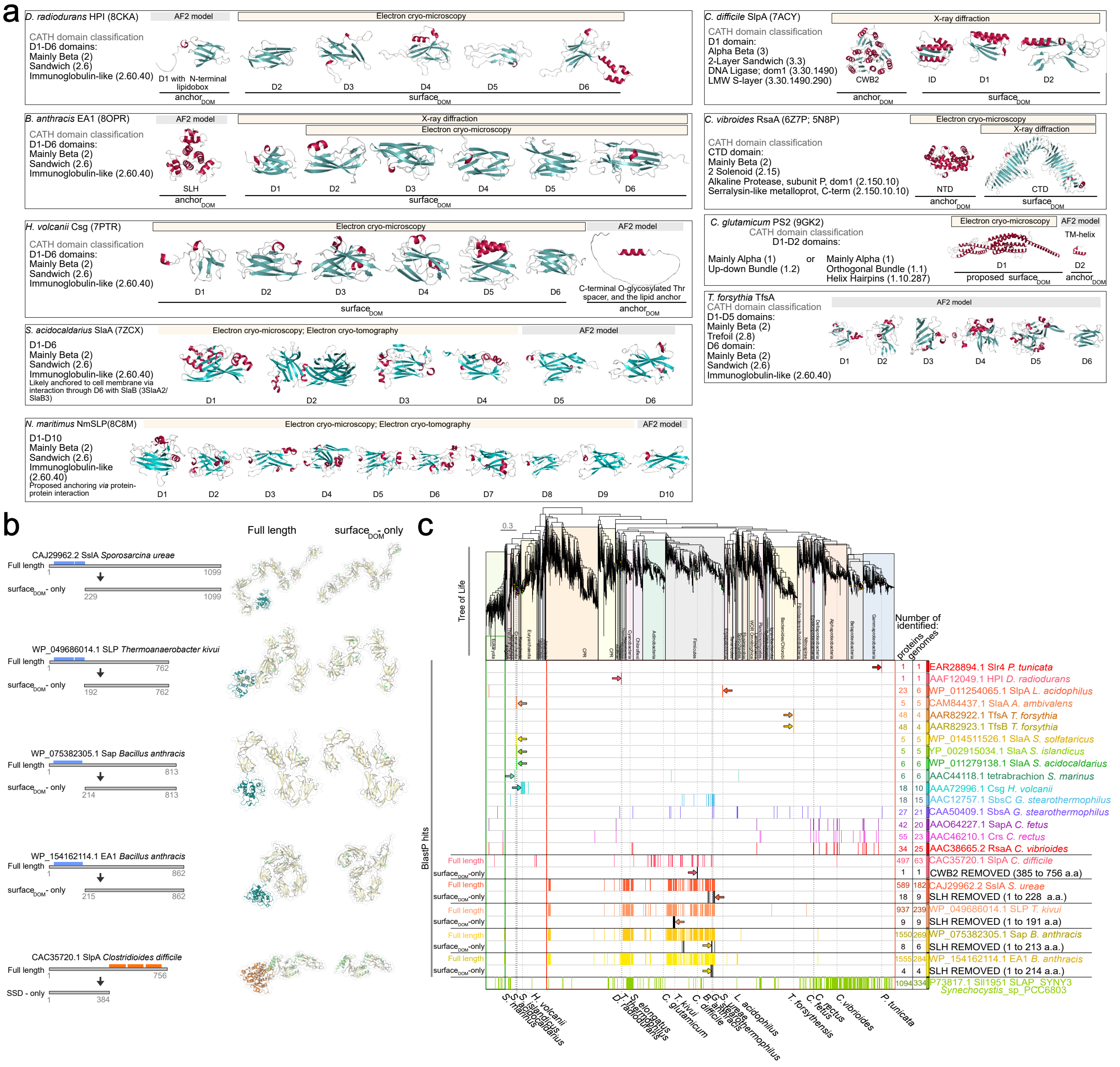


**Supplementary Fig. 1: Common anchor$_{DOM}$s mask lineage-specific distribution of SLLP surface$_{DOM}$s.**

**a,** Domain organisation of representative SLLPs with known structures (HPI, EA1, Csg, SlpA, SlaA, NmSLP, PS2 and RsaA) and a single example of a distinct fold architecture awaiting experimental verification (TfsA). CATH domain classification is that of the top hits from Foldseek searches. Surface$_{DOM}$s of two SLLPs from distantly related bacteria, *D. radiodurans* and *B. anthracis*, and one from archaeon *H. volcanii*, all share similar structural organisation with six mainly β-sandwich, immunoglobulin-like (Ig-L, CATH 2.60.40) domains (left panel). Ig-L domains are commonly found across the Tree of Life (Fig. 1a, and Supplementary Fig. 2b) and are the most common fold in currently known SLLPs. All three known 6xIg-L SLLPs have different anchor$_{DOM}$s representing distinct cell surface anchoring mechanisms. In addition, structures of two other Ig-L SLLPs from Archaea contain either ten domains (*N. maritimus*) or an additional Ig-L within D2 (*S. acidocaldarius*), illustrating the modularity of Ig-L surface$_{DOM}$ subdomain organisation. **b,** To account for the observed wide distribution of the CWB2 and SLH anchor$_{DOM}$s (Fig. 1a, and Supplementary Fig. 2b) commonly found in multiple proteins from a single genome (Supplementary Fig. 5), both full-length sequences and anchor$_{DOM}$-removed sequences containing only more SLLP-specific surface$_{DOM}$ (surface$_{DOM}$-only) were used in protein sequence homology searches (**c**). The correct removal of the amino acid sequence of the anchor$_{DOM}$s was verified with Interproscan and Alphafold2 structural predictions.
**c,** Mapping of the SLLP sequence homologues identified with BlastP (E-value < e-5) in 2,705 Bacteria, 315 Archaea, and 149 Eukaryota sampled across the Tree of Life (Fig. 1a). Numbers represent the total number of the identified BlastP hits (proteins) or hit-containing genome assemblies (genome). Results of the BlastP searches using surface$_{DOM}$-only query sequences (black) following the common anchor$_{DOM}$ sequence removal are presented below those of full-length sequence search results (colour-coded). A decrease in the numbers of the identified genomes and proteins, and a narrower taxonomic range of the identified homologues of surface$_{DOM}$-only queries, indicated that the presence of common anchor$_{DOM}$s can mask the distinct lineage-specific distribution patterns of the surface$_{DOM}$s.

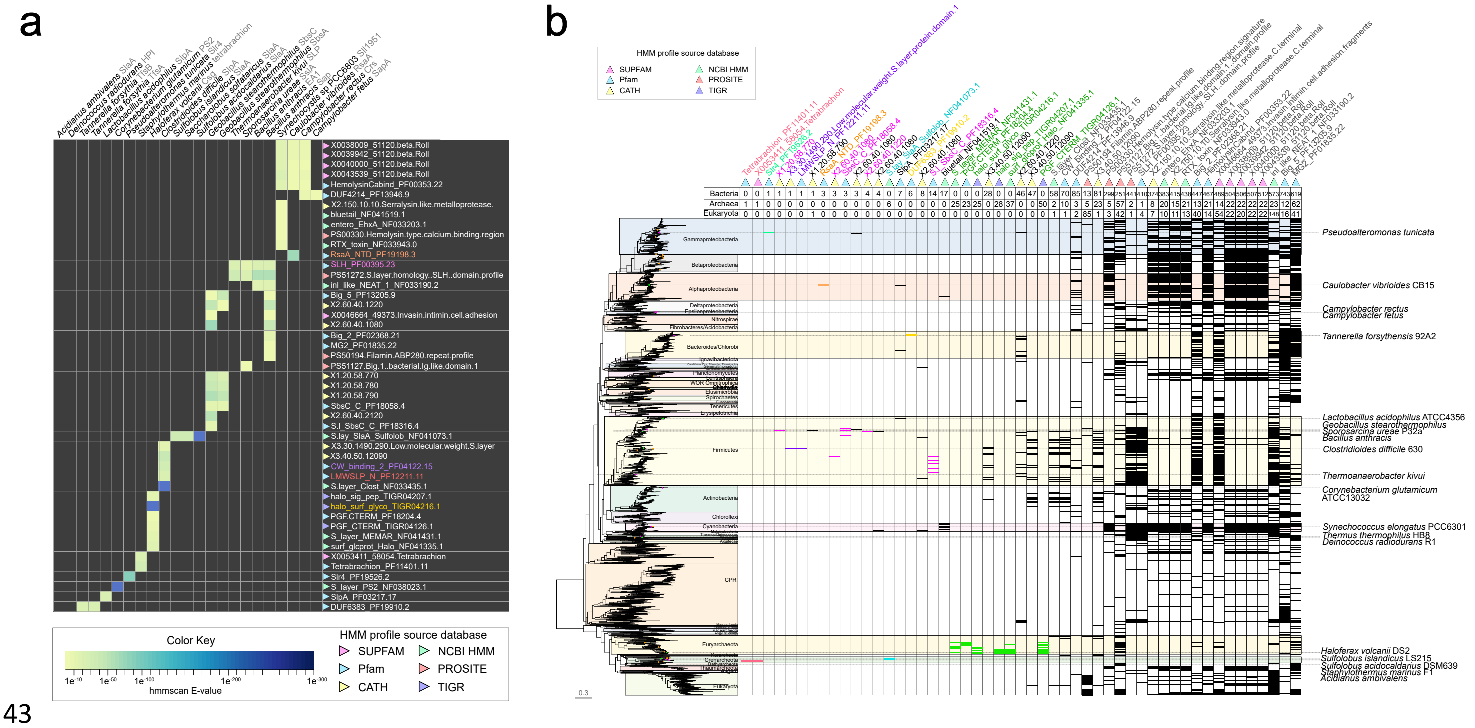


**Supplementary Fig. 2: Identification of common and unique SLLP sequence signatures.**

**a,** Mapping of the HMM profiles identified for each of the experimentally verified SLLP sequences in protein profile databases using hmmscan with search results E-values represented as a heatmap (yellow-blue). The majority of known SLLPs contain specific sequence profiles not overlapping with any other SLLPs, with overlap identified only in the cases of homologous sequences from closely related species and more conserved (**b**) β-roll profiles.

**b,** Tree of Life mapping of the sequences detected in 2,705 Bacteria, 315 Archaea, and 149 Eukaryota genomes using hmmsearch with HMM sequence profiles identified in verified SLLPs (**a**), with number of genomes encoding detected sequences above the mapping. Sequences identified with HMM profiles were found in genomes from all three domains of life (grey), more narrowly in multiple phyla (black), or within a single phylum (colour-coded). The more common profiles included immunoglobulin-like, and β-roll folds; calcium-binding motifs; and SLH and CWB2 cell wall anchoring domains. The more narrowly distributed profiles are mostly annotated in databases as specific to SLLPs.

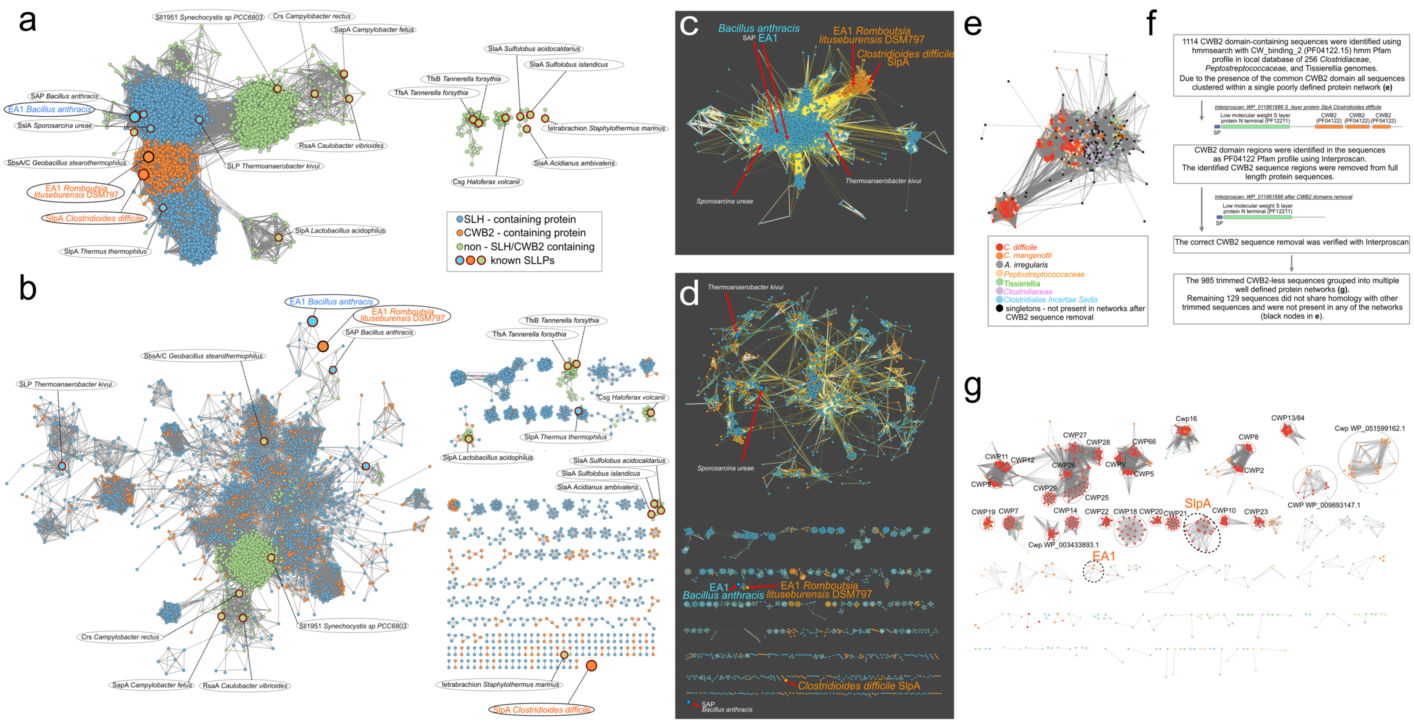


**Supplementary Fig. 3: The presence of common and swappable anchor$_{DOM}$s can cause an incorrect homology inference for different SLLPs and S-layer-associated proteins.**

**a-g,** Protein similarity networks of full-length sequences (**a, c, e**) or sequences after removal of common SLH and CWB2 domains (**b, d, g**). The networks of proteins sampled across the Tree of Life (**a** and **b**) were constructed using homologues of all SLLPs identified using BlastP searches in 2,705 Bacteria, 315 Archaea, and 149 Eukaryota (Supplementary Fig. 1c). The Firmicutes protein networks (**c** and **d**) were constructed with all CWB2 and SLH-containing proteins identified with hmmsearch in 828 Firmicutes genome assemblies.

**f,** An example of the workflow used in the analysis of the *Clostridiaceae-Tissierellia-Peptostreptococcaceae* (CTP) CWB2-protein networks (**e**-**g**). In the full-length sequence analyses, CWB2 (orange) and SLH-proteins (blue) are grouped within distinct clusters (**a** and **c**). Following the removal of these common cell wall anchoring domains, the remaining functional domains of the cell wall-anchored proteins were separated into multiple smaller networks (**b** and **d**). *R. lituseburensis* EA1 initially grouped with *C. difficile* SlpA as they share the common CWB2 anchor$_{DOM}$, however in the functional domain networks, CWB2-EA1 grouped with *B. anthracis* SLH-EA1, with which it shares the surface$_{DOM}$ fold, indicating likely SLH/CWB2 anchor swap and suggesting possible acquisition *via* LGT. This demonstrates how the presence of common anchoring domains can mask the sequence homology and evolution of the often less conserved surface$_{DOM}$ and other functional domains. This was particularly stark within the CTP networks (**e** and **g**), where all of the often functionally and structurally unrelated S-layer-associated and cell wall-binding CWB2-proteins clustered within a single 'hairball' in full-length sequence analyses (**e**), but were separated in the functional domain-only analyses (**g**). In the Tree of Life (**b**) and Firmicutes (**d**) analyses, these functional domain networks often (14 networks in **b**, and 33 in **d**) contained sequences of SLH and CWB2 proteins, consistent with common SLH/CWB2 anchoring domain swaps (Fig. 4c-f). The majority of the non-SLH/CWB2 proteins (green) in the Tree of Life networks (**a** and **b**) clustered together and included SLLPs from *Synechocystis, Caulobacter,* and *Campylobacter,* all of which contain a common calcium-binding motif and β-roll domains (Supplementary Fig. 2).

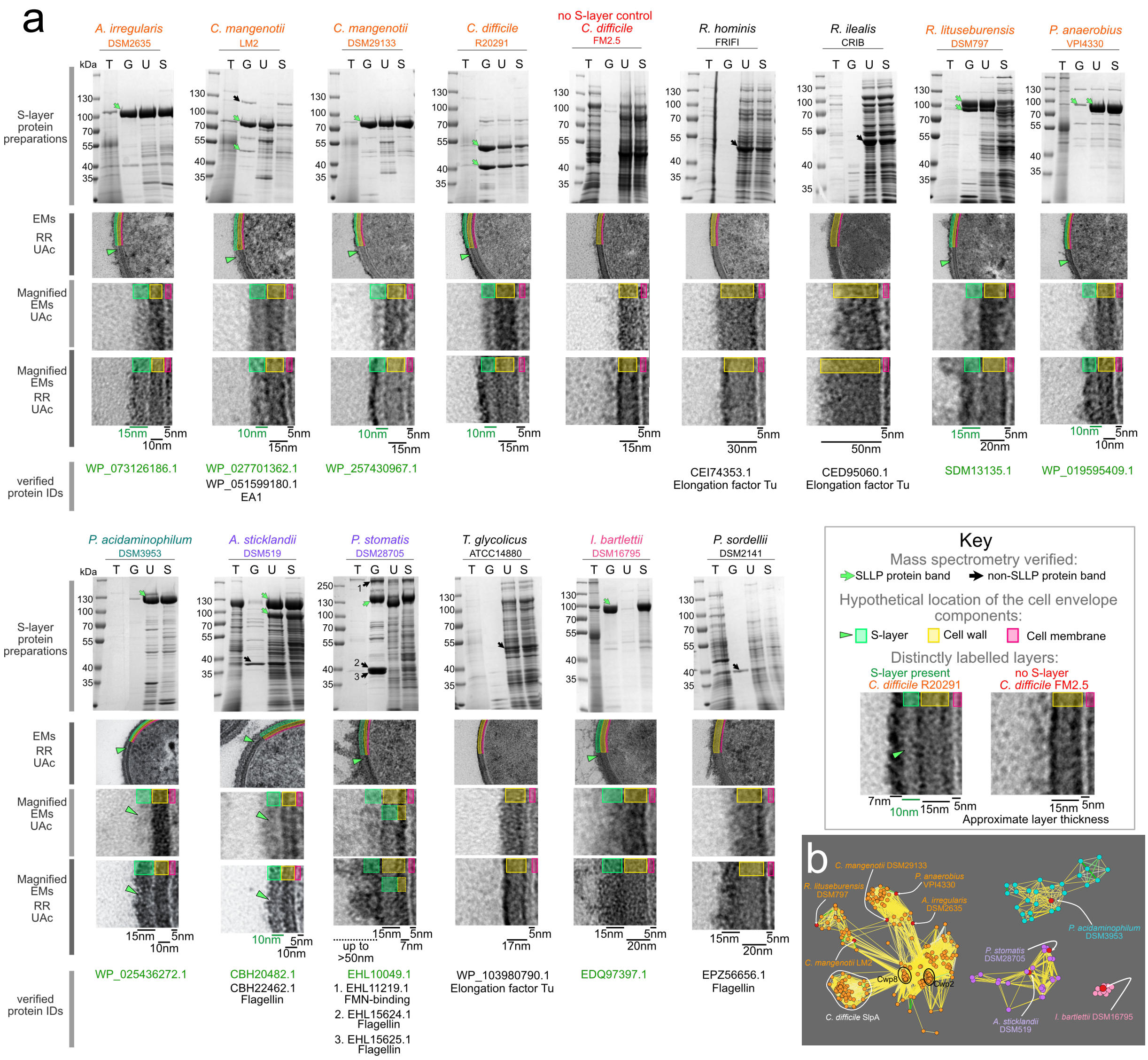


**Supplementary Fig. 4: Investigating the presence of S-layers and SLLPs in *Peptostreptococcaceae*.**

**a,** SDS-polyacrylamide gels of S-layer preparations isolated using low pH glycine (G), urea (U), or SDS (S). Proteins present in the spent culture media were precipitated with trichloroacetic acid (T). The molecular weight ladder is shown (kDa). NCBI identifiers (verified protein IDs) refer to the SLLP (green arrows) and non-SLLP (black arrows) protein bands analysed using mass spectrometry. Transmission electron micrographs of *Peptostreptococcaceae* cell sections (EMs) and magnified regions of the cell surface (Magnified EMs) were stained with uranyl acetate (UAc) to visualise apparent layers of varying electron density on the cell surface and ruthenium red (RR) to visualise cell surface glycopolymers. Scale bars indicate approximate thickness of the layers most likely corresponding to the S-layer (green), and the cell wall (yellow). The approximate location of the cell membrane was highlighted as the electron-dense edge of the cytosol (pink). Two *Romboutsia* isolates (*R. hominis* FRIFI and *R. ilealis* CRIB) with no apparent evidence of the S-layer in the EMs and SDS-PAGE gels, displayed much thicker cell wall layers, which could reflect the adaptation of these bacteria to the possible evolutionary loss of the S-layer or at least its absence in the tested experimental conditions. In *C. difficile* FM2.5, the control strain for absence of the S-layer, the presence of high-intensity bands in the spent media and in the SDS and urea extractions most likely reflects increased cell susceptibility to these chemical agents and autolysis of the S-layer null strain. Evidence of cell lysis was also observed in chemically treated species with no evidence of S-layer (*R. hominis* FRIFI, *R. ilealis* CRIB, and *T. glycolicus*) where the highest intensity bands correspond to homologues of a highly abundant cytosolic elongation factor protein. Interpretation of the EMs was less clear in species such as *P. stomatis* which displays differential staining between the UAc and RR+UAc samples, consistent with particularly strong surface glycosylation. This species also exhibits a high number of flagella, apparent in the micrographs and as flagellin protein bands.

**b**, EGN protein similarity networks generated using the experimentally identified SLLP sequences (red) and their homologues identified across 453 *Peptostreptococcaceae*. Each network represents a distinct surface$_{DOM}$ fold and anchoring mechanism (colour-coded as in Fig. 2b and 4a), demonstrating that the identified distinct S-layer types do not share any sequence homology.

a

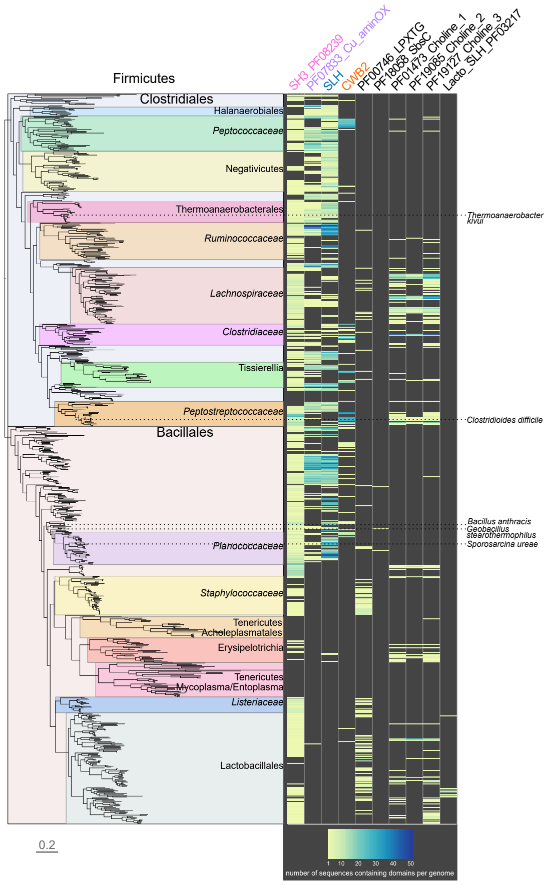


b

Number of hmmsearch hits against 828 Firmicutes genomes

| HMM profile | total proteins | mean proteins per genome | total genomes | % total genomes |
|---|---|---|---|---|
| PF08239 SH3 | 1775 | 3.1 | 581 | 70.2 |
| PF00395 SLH | 4039 | 12.5 | 322 | 38.9 |
| PF07833 Cu_aminOX | 2398 | 9.0 | 265 | 32.0 |
| PF19127 Choline_bind_3 | 845 | 6.1 | 139 | 16.8 |
| PF01473 Choline_bind_1 | 591 | 5.4 | 110 | 13.3 |
| PF00746 LPXTG | 236 | 2.2 | 105 | 12.7 |
| PF04122 CWB2 | 773 | 9.8 | 79 | 9.5 |
| PF19085 Choline_bind_2 | 293 | 5.4 | 54 | 6.5 |
| PF03217 Lacto_SLH | 41 | 5.1 | 8 | 1.0 |
| PF18058 SbsC | 10 | 2.5 | 4 | 0.5 |

c

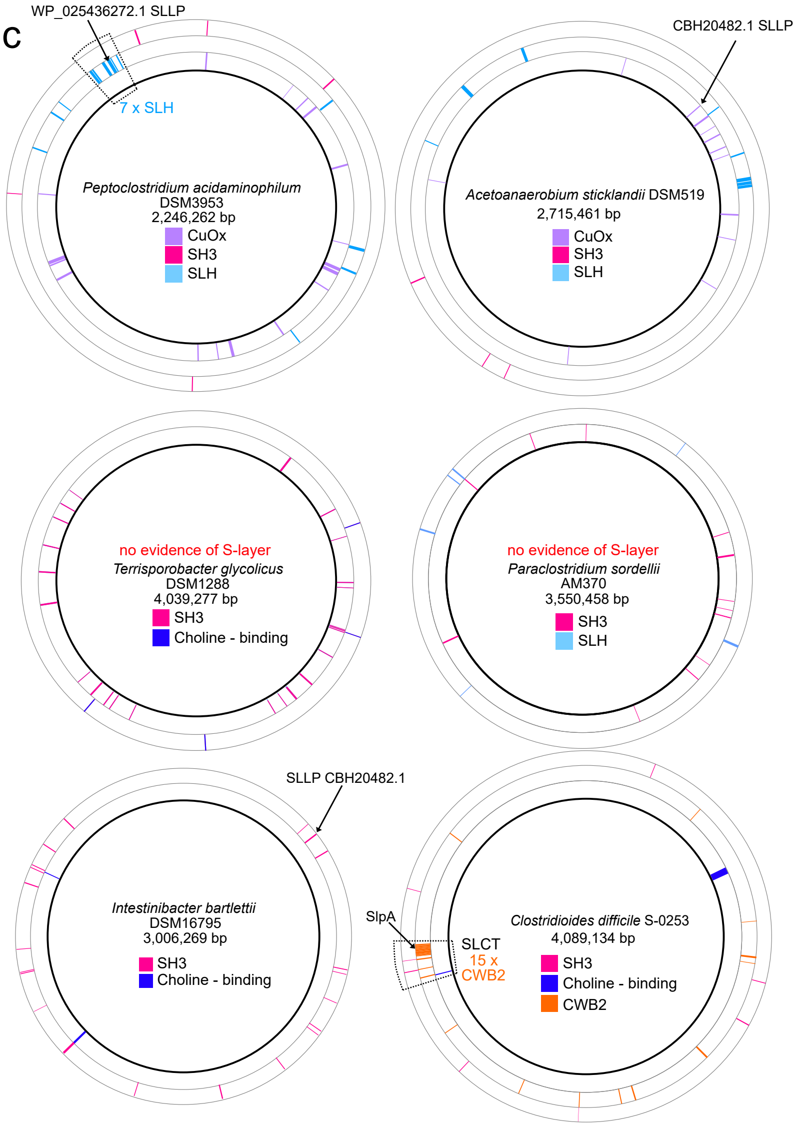


**Supplementary Fig. 5: Distribution of the different cell surface anchoring domains found across Firmicutes.**

**a,** Species tree mapping of the number (yellow-blue heatmap) of proteins containing the investigated cell surface anchoring domains detected with hmmsearch in 828 Firmicutes genomes.

**b,** Numbers of proteins and genomes containing the investigated $anchor_{DOM}$s in (**a**).

**c,** Genome location of the genes encoding $anchor_{DOM}$-containing proteins mapped onto the circular representations of the available closed genomes of the *Peptostreptococcace* species experimentally investigated in this study. The black arrows indicate the location of the genes encoding the identified SLLPs. Regions containing higher numbers of $anchor_{DOM}$-protein coding genes were outlined (dashed lines).

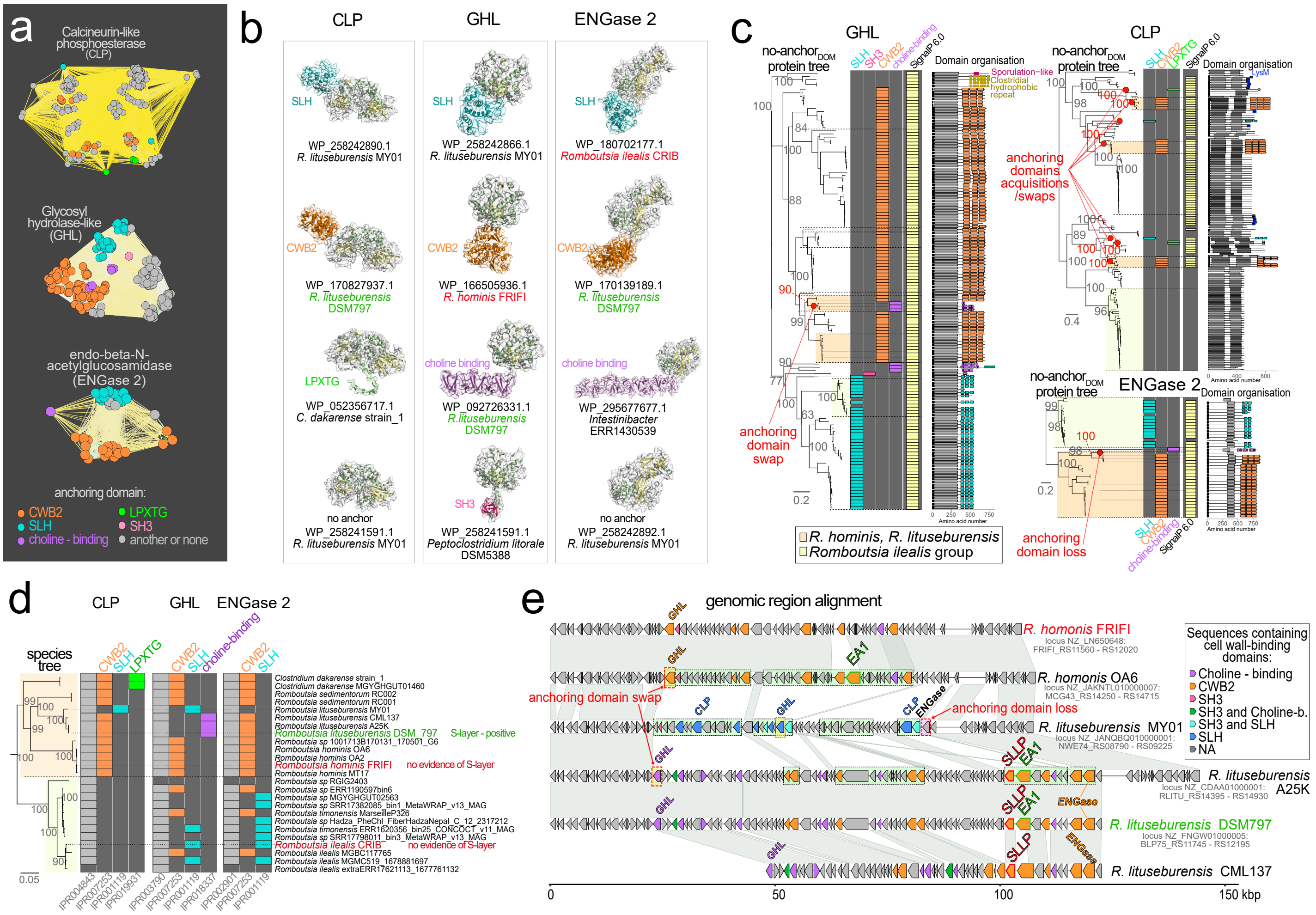


**Supplementary Fig. 6 Modular architecture and evolution of the S-layer and cell wall-anchoring systems in *Romboutsia.***

**a,** Protein similarity networks with well-supported evidence of cell wall anchor switching, gain and/or loss identified in the *Peptostreptococcaceae*.

**b,** AlphaFold2 structure prediction of the representative proteins from (a-c) containing alternative anchor$_{DOM}$ (colour-coded as in a) or no anchor attached to the same functional domain (no anchor).

**c,** Protein phylogenies of homologues from (a) mapped with the identity of their respective anchor$_{DOM}$s. Clear patterns of anchor$_{DOM}$ loss, acquisition, and swap were annotated on the trees (red circles). The presence of secretion signals (pale yellow), detected with SignalP 6.0, suggested that none of the indicated changes were associated with re-localisation to the cytosol. Protein domain architecture identified with Interproscan (Domain organisation) is presented for each homologue with conserved 'functional domains' (grey), signal peptides (black), and anchor$_{DOM}$s (colour-coded).

**d,** Presence of the attached functional domains (grey) and different anchor$_{DOM}$s (orange CWB2, cyan SLH, green LPXTG, magenta choline-binding) in the *Romboutsia* homologues from (a) were mapped onto the *Romboutsia* species tree.

**e,** Alignment of *R. lituseburensis* and *R. hominis* genomic regions demonstrating replacement of CWB2 protein-containing region with SLH region in *Romboutsia lituseburensis* MY01 (dashed outlines), a single anchor$_{DOM}$ swap and a single loss in individual genes (red arrows). SLLP is the main S-layer protein homologous to *C. difficile* SlpA identified in S-layer preparations from *R. lituseburensis* DSM797 (Supplementary Fig. 4a). EA1 is the homologue of *Bacillus* SLLP (EA1), with SLH domain-swapped for CWB2 (Fig. 4e).

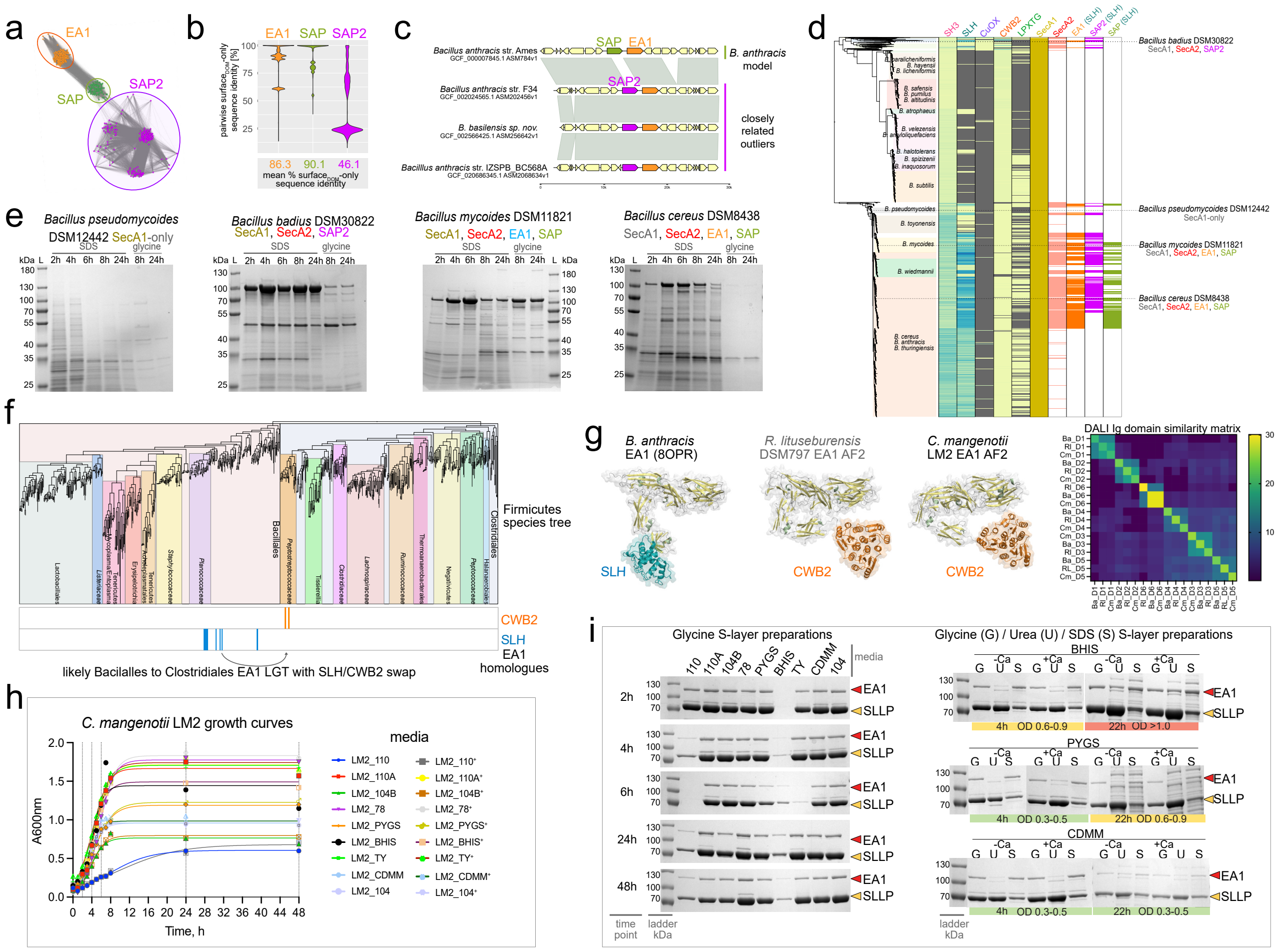


**Supplementary Fig. 7: Evolution of EA1/SAP S-layer proteins in *Bacillus* and *Peptostreptococcaceae*.**

**a,** Protein homology network analysis of a non-redundant set of SLLP homologues identified across all available (7,377) *Bacillus* genome assemblies revealed a third distinct SAP/EA1 family member (SAP2) which includes a recently identified *B. cereus* S-layer protein SL2[52].

**b,** Pairwise sequence identity comparisons of surface$_{DOM}$s indicate that SAP and EA1 are more conserved, while SAP2 provides higher SLLP diversity, reminiscent of SlpA in *C. difficile* (Fig. 5d).

**c,** Genome alignment of closest SAP2-containing relatives of *B. anthracis* Ames, indicates that SAP2 and SAP are interchangeable.

**d**, Mapping of all three family members across the *Bacillus* phylogeny revealed a differential distribution, similar to that observed in *C. difficile* (Fig. 5b), providing evidence of frequent SLLP lateral gene transfers across *Bacillus*. Similarly to *Peptostreptococcaceae* (Fig. 4c), SLH-anchored SLLPs codistribute with SecA2 secretion system. Interestingly, a large group within *Bacillus cereus/anthracis/thuringiensis* contained no genome-encoded SLLP and SecA2 homologues, expanded SH3- or reduced SLH-protein repertoires.

**e,** SDS protein gels of glycine and SDS S-layer extracts sampled from across the growth cycle of different *Bacillus* species (**d**). The single species with no identifiable SLLP and SecA2 homologues (*Bacillus pseudomycoides* DSM12442) displayed no intense protein bands, consistent with possible S-layer loss.

**f,** Mapping CWB2- and SLH-containing EA1 homologues identified in the network analyses (Supplementary Fig. 3) onto Firmicutes species tree revealed anchoring domain (CWB2/SLH) swapping and lateral gene transfer (LGT) between members of *Bacillaceae* and *Peptostreptococcaceae.*

**g,** Comparison of the determined *B. anthracis* SLH-EA1 structure with AlphaFold2 models of *R. lituseburensis* and *C. mangenotii* CWB2-EA1. The immunoglobulin-like EA1 surface$_{DOM}$ subdomains (Supplementary Fig. 1a) were compared with pairwise Distance matrix alignments (DALI) of their 3D structures (yellow-blue heatmap). The structural comparisons are consistent with the conservation of the surface$_{DOM}$ and SLH/CWB2 swap of the anchor$_{DOM}$.

**h-i** Presence of the EA1 in *C. mangenotii* LM2 S-layer preparations (**i**) was tested across different growth phases (**h**) in different culture conditions (media) with (+Ca) and without (-Ca) calcium supplementation. S-layers were extracted using three different chemical methods (glycine, G; urea, U; and SDS, S). In SDS-polyacrylamide gels of S-layer protein preparations (**i**), *C. mangenotii* CWB2-EA1 (red arrowhead) appears to be a minor S-layer associated protein, alongside the SlpA-type CWB2-SLLP (yellow arrowhead). Identification of CWB2-EA1 in *C. mangenotii* S-layer preparations indicates that the anchor$_{DOM}$-swapped EA1 is functional.

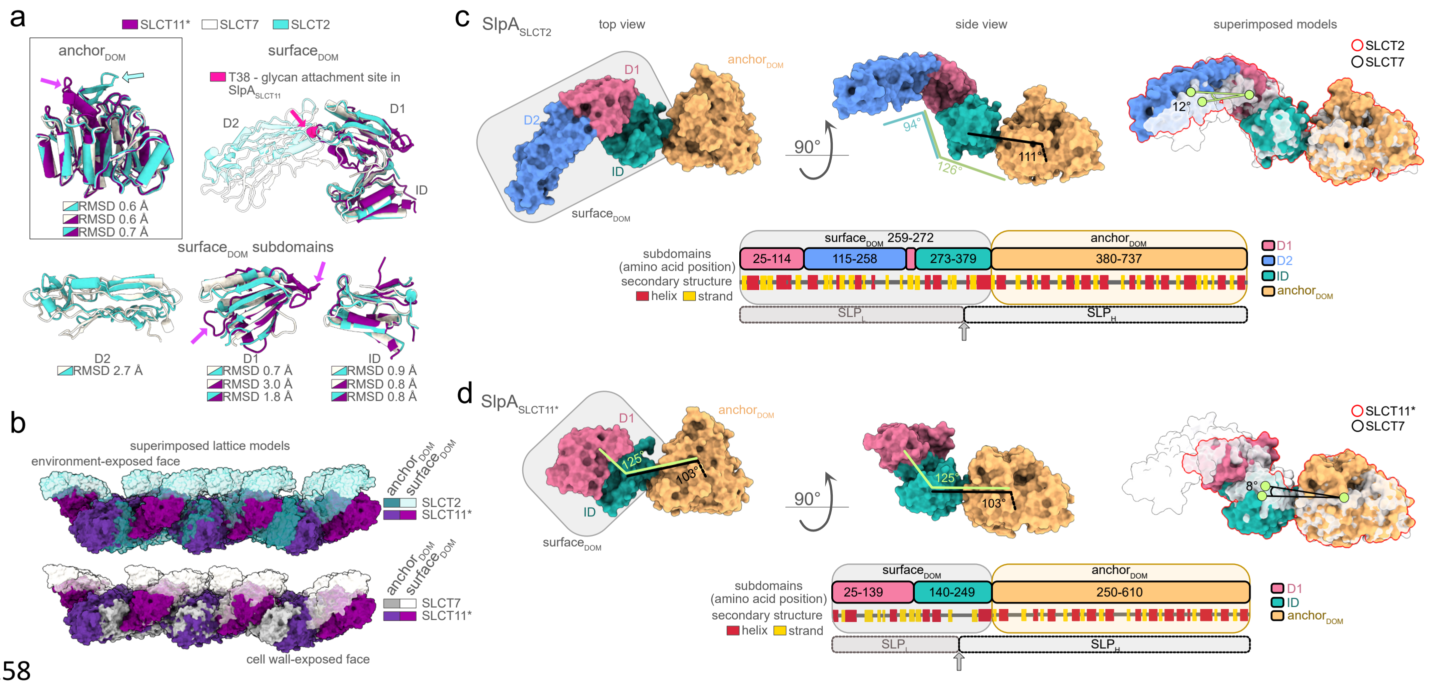


**Supplementary Fig. 8: Different architectures of *C. difficile* SlpA crystal lattices.**

**a,** Structural alignments of anchor$_{DOM}$s (top left), surface$_{DOM}$s (top right), and individual surface$_{DOM}$ subdomains (bottom) from all three experimentally determined SlpA$_{SLCTs}$ (SlpA$_{SLCT2}$, cyan; SlpA$_{SLCT7}$, white; SlpA$_{SLCT11}$; magenta). Calculated structural differences between polypeptide backbones of each SlpA$_{SLCT}$ are represented as root mean square deviation (RMSD) values. Within the more conserved anchor$_{DOM}$s, the additional β-hairpin of SlpA$_{SLCT2}$ (cyan arrow) is located on the outer anchor$_{DOM}$ face, between surface$_{DOM}$s, and the additional α-helix and 3,10-helix of SlpA$_{SLCT11}$ (magenta arrow) are at the surface-exposed face of the lattice. The most extensive differences were observed within surface$_{DOM}$ subdomains D2 and D1. The extended D2, which contains several flexible loops in SlpA$_{SLCT2}$ and SlpA$_{SLCT7}$, is absent from SlpA$_{SLCT11}$. D1 of SlpA$_{SLCT11}$ contains three shorter α-helices packed against a 4-strand mixed β-sheet, as opposed to two longer α-helices against a 5-strand mixed β-sheet in SlpA$_{SLCT7}$ and SlpA$_{SLCT2}$. Only SlpA$_{SLCT11}$ contains additional extended loops in D1 (magenta arrows). As opposed to other studied SlpA$_{SLCTs}$, SlpA$_{SLCT11}$ from the wild-type *C. difficile* Ox247 strain is heavily glycosylated and surface$_{DOM}$ threonine 38 (pink sphere and arrow) has a known *O*-linkage to a complex glycan[1].

**b,** Superimposed surface representation of SlpA$_{SLCT2}$ and SlpA$_{SLCT11}$ (top), or SlpA$_{SLCT7}$ and SlpA$_{SLCT11}$ (bottom) crystal arrays. The most striking differences are apparent in the relative position of surface$_{DOM}$ from SlpA$_{SLCT11}$ (purple), compared to that from SlpA$_{SLCT2}$ (pale cyan, top) and SlpA$_{SLCT7}$ (white, bottom). Importantly, native heavy glycosylation of SlpA$_{SLCT11}$ prevented all attempts of *in vitro* crystallisation, and only SlpA$_{SLCT11}$ preparation from glycosylation-deficient mutant strain Δ*orf2* (denoted with an asterisk) was amenable to X-ray crystallography. Whether native glycosylation can structurally compensate for the lack of the subdomain D2 awaits experimental verification, however, our experiments demonstrated that the SlpA$_{SLCT11}$ from Δ*orf2* can crystalise *in vivo* and form S-layers on the Δ*orf2* cell surface, and re-crystalise *in vitro* following glycine extraction.

**c,d**, Surface representation of SlpA$_{SLCT2}$ (**c**) and SlpA$_{SLCT11}$ (**d**) coloured by subdomains of surface$_{DOM}$ (interacting domain, ID, teal; D1, pink; D2, blue), and anchor$_{DOM}$ (orange), as viewed from the external environment (left) and side (middle). The schematics (bottom of each panel) depict domain organisation (colour-coded), secondary structure content (yellow and red rectangles), and the position of the post-translational peptidase cleavage site (grey arrow) generating two mature polypeptides (SLP$_L$ and SLP$_H$). Structures of SlpA determined in this study (right, coloured surface with red overlay) were superimposed onto SlpA$_{SLCT7}$ structure (7ACY; white, semi-transparent surface). Angles measured between centres of mass of each subdomain (ID, D1, D2) or anchor$_{DOM}$ show that a common element of surface$_{DOM}$ (D1-ID) assumes a similar orientation (125-126°) relative to the anchor$_{DOM}$ across the determined SlpA$_{SLCT}$ structures. Compared to SlpA$_{SLCT7}$, D2 in SlpA$_{SLCT2}$ undergoes a 12° rotation relative to the D1 centre of mass (**c**), placing it further away from the anchor$_{DOM}$ array plane and widening the span of interactions across the surface$_{DOM}$s within the crystal lattice (Supplementary Fig. 9). Additionally, a specific architecture of SlpA$_{SLCT11}$ lattice (**d**) is impacted by an 8° rotation of the ID relative to the anchor$_{DOM}$ centre of mass, bringing the surface$_{DOM}$ towards the cell wall, when compared to SlpA$_{SLCT7}$ and SlpA$_{SLCT2}$ arrays, therefore altering the packing of the anchor$_{DOM}$s (Fig. 5e) and contributing to interactions at a lower plane of the lattice (Supplementary Fig. 9).

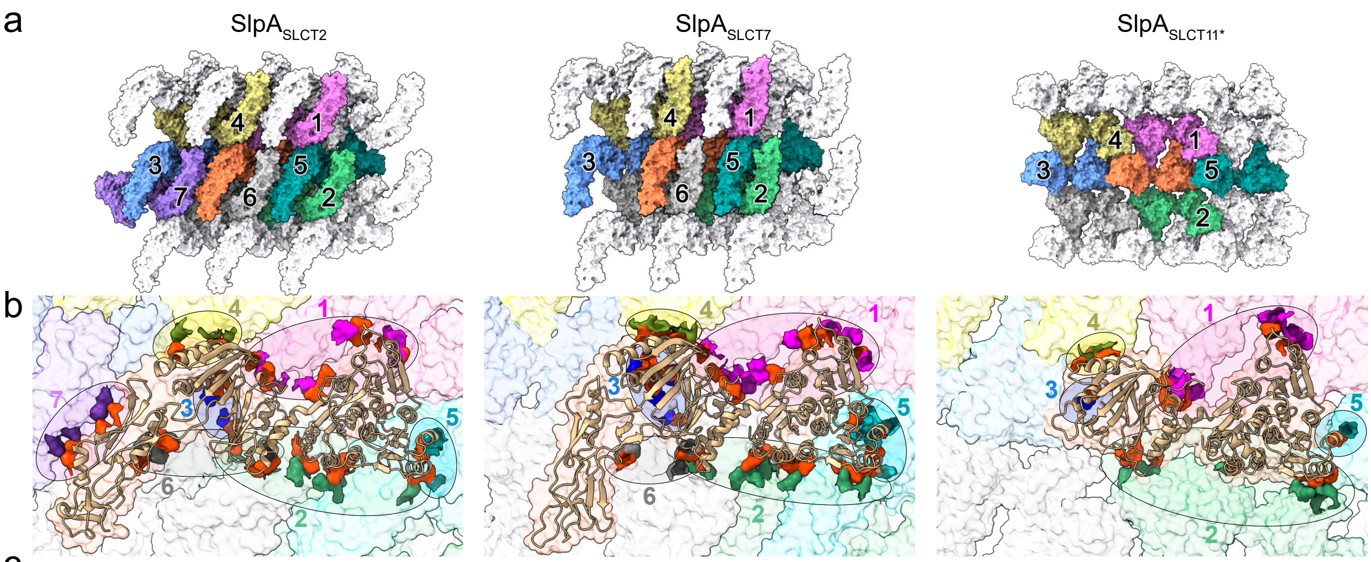


c

| Interaction type | Interacting domains | | | |
|---|---|---|---|---|
| | AD:AD | AD:SD | SD:SD | total |
| H-Bond | 8 | 16 | 11 | 35 |
| Salt Bridge | 3 | 6 | 6 | 15 |
| total | 11 (22 %) | 22 (44 %) | 17 (34 %) | 50 (100%) |

| Interaction type | Interacting domains | | | |
|---|---|---|---|---|
| | AD:AD | AD:SD | SD:SD | total |
| H-Bond | 9 | 21 | 7 | 37 |
| Salt Bridge | 6 | 8 | 1 | 15 |
| total | 15 (29 %) | 29 (56 %) | 8 (15 %) | 52 (100%) |

| Interaction type | Interacting domains | | | |
|---|---|---|---|---|
| | AD:AD | AD:SD | SD:SD | total |
| H-Bond | 2 | 17 | 4 | 23 |
| Salt Bridge | - | 4 | - | 4 |
| total | 2 (7 %) | 21 (78 %) | 4 (15 %) | 27 (100%) |

d

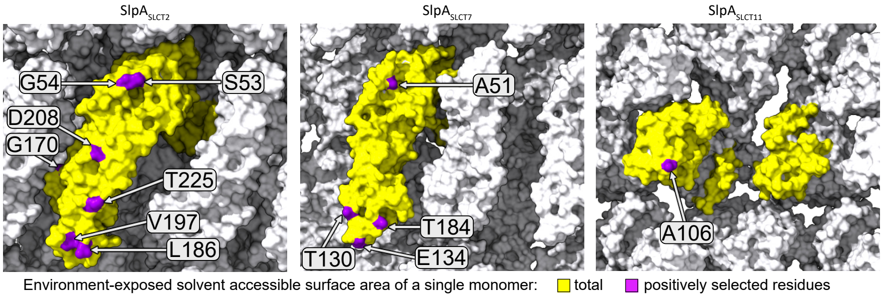


Environment-exposed solvent accessible surface area, Å$^2$

| Crystal lattice | $SlpA_{SLCT2}$ | $SlpA_{SLCT7}$ | $SlpA_{SLCT11}$ |
|---|---|---|---|
| Total exposed area of a single monomer | 16288.0 | 16115.8 | 14058.6 |
| Average exposed area of a single residue [lowest - highest values] (% total) | 62.2 ± 41.5 [4.3 - 187.5] (0.03 - 1.15 %) | 63.2 ± 41.9 [4.5 - 188.8] (0.03 - 1.17 %) | 73.2 ± 49.5 [5.2 - 252.5] (0.04 - 1.80 %) |
| Exposed area of all positively selected residues | 511.8 | 392.7 | 89 |
| Average exposed area of a positively selected residue [lowest - highest values] (% total) | 73.1 ± 39.7 [38.3 - 154.2] (0.24 - 0.95 %) | 78.5 ± 44.8 [22.9 - 136.8] (0.14 - 0.85 %) | 89 [single residue] (0.63 %) |

**Supplementary Fig. 9: Both $surface_{DOM}$ and $anchor_{DOM}$ are involved in *C. difficile* SlpA crystal lattice formation.**

**a,** Surface representation of $SlpA_{SLCT2}$, $SlpA_{SLCT7}$ and $SlpA_{SLCT11}$ crystal lattices (13, 12 and 15 monomers shown, respectively). A single SlpA molecule (orange) establishes direct crystal contacts with neighbouring molecules (numbered and coloured).
**b,** Zoom view of lattices from **a**, centred on a single SlpA molecule (cartoon representation), with each of the neighbouring units depicted as a semi-transparent lattice surface (coloured, and numbered as per morphological unit in **a**). Each of the interaction interface regions between the central SlpA molecule and its neighbours is outlined with ellipses. Residues in direct crystal contact between the central molecule (orange) and neighbouring molecules (coloured accordingly) were depicted as dark, opaque surfaces. Despite the striking differences in lattice architecture between $SlpA_{SLCT11}$ and the other two SlpAs, all three arrays include a common set of five interaction interfaces comprising the central unit and its neighbouring molecules 1-5. The presence of D2 and a more protruding position of the ID (Supplementary Fig. 8c) in $SlpA_{SLCT2}$ and $SlpA_{SLCT7}$ contribute to interactions that build an additional interface (6), not present in $SlpA_{SLCT11}$. Furthermore, the rotation of D2 relative to D1 in $SlpA_{SLCT2}$ compared to $SlpA_{SLCT7}$ (Supplementary Fig. 8b) allows additional contacts (interface 7) with a seventh molecule, generating a more tightly packed environment-exposed lattice surface.
**c,** In all $SlpA_{SLCT}$s, the majority of inter-molecular interactions were found between S-layer $surface_{DOM}$ (SD) and $anchor_{DOM}$ (AD). The number of hydrogen bond and salt bridge interactions identified between the central molecule and neighbouring molecules are reflective of different packing and crystal lattice organisation. This was particularly apparent for the $surface_{DOM}$:$surface_{DOM}$ interfaces where a higher number of interactions represents tighter $surface_{DOM}$ packing, and a higher number of molecules in direct contact (6 in $SlpA_{SLCT11}$, 7 in $SlpA_{SLCT7}$, or 8 in $SlpA_{SLCT2}$). Interestingly, the relative proportion of H-bonds (H-b) to salt bridges (Sb) at the lattice interfaces, was higher in $SlpA_{SLCT11}$ (H-b/Sb = 5.75) compared to $SlpA_{SLCT7}$ (H-b/Sb = 2.47) and $SlpA_{SLCT2}$ (H-b/Sb = 2.33).
**d,** Environment-exposed surface plane of the $SlpA_{SLCT}$ crystal lattice (white surface) was manually selected, and a solvent-exposed surface area (SASA) was estimated for a single molecule (yellow surface) within each lattice, using Chimera. Cumulative and average (+/-standard deviation) solvent exposure was calculated (summarised in a table, right panel) for the surface residues of the environment-exposed surface plane. A substitution of a single positively selected amino acid (purple surface) is estimated to affect at most 0.95% of the entire S-layer cell surface, in a hypothetical scenario of a homogenous SlpA lattice covering the entire *C. difficile* cell.

**a**

| | protein | Number of uniqe nucleotide sequences | Nucleotide alignment length (variable sites) | Mean nucleotide sequence identity | PAML: CODEML M1a vs M2a positive selection test p-value ($\chi^2$) | PAML: identified specific sites under positive selection Number P>99% (P>95%) | PAML: identified specific sites under positive selection lowest ω (dN/dS) | GARD: recombination detection inferred breakpoints | FUBAR: identified specific sites posterior probability > 90% (> 99%) positive selection | FUBAR: identified specific sites posterior probability > 90% (> 99%) negative selection | BUSTED: support for episodic diversifying selection (p-value) | MEME: number of sites with episodic positive/diversifying selection (p-value < 0.1) | Evidence of positive selection: number of sites detected in all three (two) methods | protein |
|---|---|---|---|---|---|---|---|---|---|---|---|---|---|---|
| SlpA | SLCT1 | 37 | 1710 (318) | 93.8 % | 1.37E-05 | 1 (6) | 5.8 | 1 | 5 (0) | 62 (4) | no (0.50) | 4 | 2 (3) | SLCT1 |
| SlpA | SLCT2 | 89 | 1914 (1065) | 83.3 % | 0 | 13 (16) | 4.1 | 1 | 2 (0) | 371 (172) | yes (0.0053) | 42 | 1 (7) | SLCT2 |
| SlpA | SLCT3 | 38 | 2220 (942) | 83.4 % | 0 | 13 (18) | 8.9 | 2 | 7 (1) | 204 (15) | yes (0.0024) | 13 | 3 (6) | SLCT3 |
| SlpA | SLCT4 | 95 | 1920 (421) | 95.2 % | 3.10E-12 | 4 (6) | 5.6 | 1 | 10 (1) | 48 (10) | yes (0.0014) | 16 | 2 (5) | SLCT4 |
| SlpA | SLCT6 | 40 | 2097 (360) | 96.2 % | 1.69E-10 | 4 (6) | 6.6 | 1 | 4 (0) | 64 (8) | yes (0.000031) | 13 | 0 (3) | SLCT6 |
| SlpA | SLCT7 | 72 | 2109 (552) | 92.6 % | 0 | 6 (13) | 6.8 | 1 | 4 (0) | 134 (14) | yes (3.7e-7) | 16 | 2 (3) | SLCT7 |
| SlpA | SLCT8 | 69 | 2100 (556) | 91.2 % | 2.06E-09 | 2 (7) | 3.1 | 1 | 7 (2) | 108 (17) | yes (0.00014) | 16 | 3 (6) | SLCT8 |
| SlpA | SLCT9 | 48 | 1827 (590) | 92.2 % | 6.66E-16 | 6 (9) | 9.4 | 1 | 12 (0) | 73 (4) | yes (0.000033) | 14 | 5 (6) | SLCT9 |
| SlpA | SLCT10 | 103 | 1551 (712) | 91.1 % | 0 | 9 (12) | 3.7 | 1 | 7 (1) | 172 (53) | yes (2.1e-13) | 31 | 4 (7) | SLCT10 |
| SlpA | SLCT11 | 26 | 1719 (492) | 94.6 % | 0.00408 | 0 (1) | 7 | 0 | 1 (0) | 195 (23) | yes (0.0084) | 6 | 0 (1) | SLCT11 |
| | Cwp2 | 228 | 1593 (531) | 96.3 % | 1 | - | - | 1 | 1 (0) | 223 (118) | no (0.50) | 9 | 0 (1) | Cwp2 |
| | Cwp8 | 205 | 1446 (336) | 98.3 % | 1 | - | - | 0 | 1 (0) | 143 (48) | yes (0.021) | 6 | 0 (0) | Cwp8 |
| | Cwp66 | 263 | 1758 (1270) | 77.1 % | 0.00956 | 0 | - | 1 | 1 (0) | 519 (381) | no (0.30) | 26 | 0 (1) | Cwp66 |

**b**

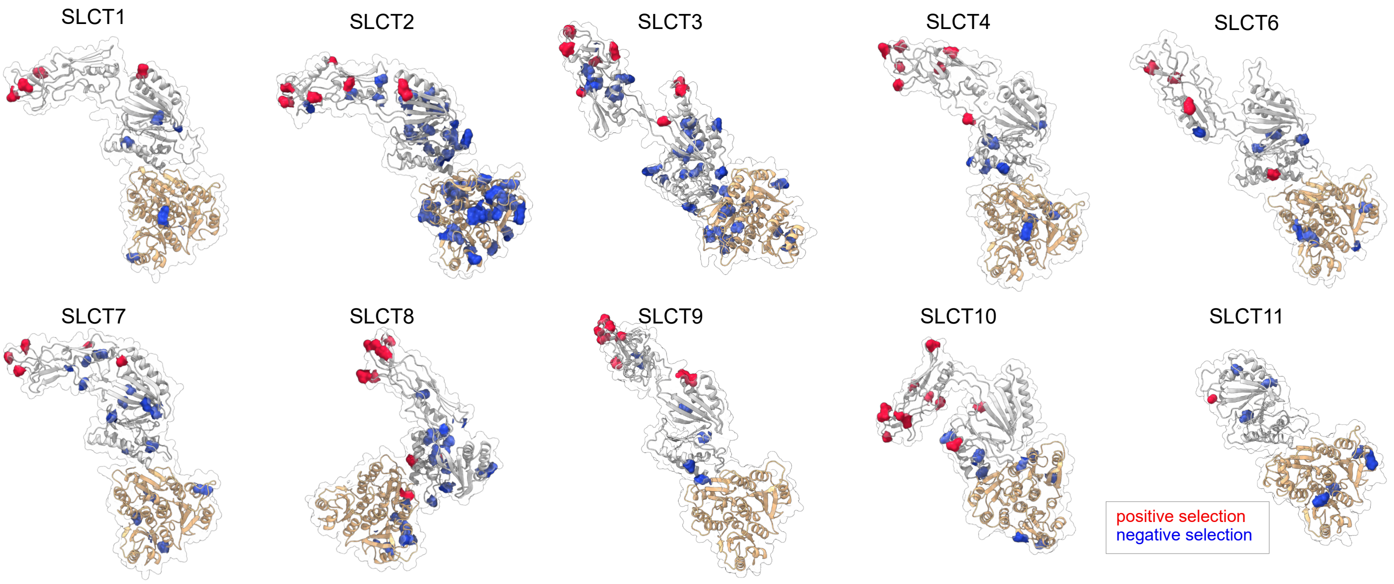


**Supplementary Fig. 10: Investigating the patterns of adaptive evolution in *C. difficile* SlpA**

**a,** Summary table of the adaptive evolution inference for *C. difficile* SLLP (SlpA), and three S-layer-associated proteins including two SlpA homologues (Cwp2 and Cwp8, Fig. 3b) and the second least conserved SCO90 Cwp66 (Fig. 5d). SlpA$_{SLCT}$s were analysed separately to enable accurate sequence alignments of the hypervariable surface$_{DOM}$s, account for possible inter-SlpA$_{SLCT}$ recombinations (Fig. 5c), and provide replicas, facilitating the investigation of trends conserved across all SlpA$_{SLCT}$s. Statistically significant evidence of positive selection was identified for all analysed SlpA$_{SLCTs}$ using all methods (CODEML M1a vs M2a, p-value < 0.01; FUBAR, posterior probability > 90%; BUSTED, p-value < 0.01; MEME, p-value < 0.1) with a single exception of no support for episodic diversifying selection in SlpA$_{SLCT1}$ (MEME, p-value = 0.5). This last observation is particularly interesting as SlpA$_{SLCT1}$ in our analyses was found mainly within clade1 (Fig. 5a,b). Only the sites identified using at least two out of three methods (PAML, FUBAR, MEME) were considered to display strong evidence of positive selection. No robust evidence of positive selection across all methods was found for the S-layer-associated proteins (Cwp2/8, Cwp66), suggesting that the observed patterns are SlpA-specific. Furthermore, Cwp2/8 and SlpA share a common CWB2-α/β-sandwich architecture, suggesting that the patterns observed for SlpA are specific to their role as SLLP and not this particular protein fold.

**b,** Cartoon representations of the SlpA$_{SLCT}$ molecules with determined structures (SLCT: 2, 7, 11) or AlphaFold2 models (SLCT: 1, 3, 4, 6, 8, 9, 10), coloured by surface$_{DOM}$ (white) and anchor$_{DOM}$ (light orange). Sites with evidence of positive selection identified using at least two different methods in **a**, were displayed as spheres and surface representations (red). All the annotated positively selected sites were on the environment-exposed face of the available experimental crystal lattices (Fig. 5f). The evidence of negative selection (blue) identified using only a single method (FUBAR), displayed no consistent distribution pattern and was found for buried residues, and surface residues from all SlpA$_{SLCT}$ faces (Fig. 5f).

**Supplementary Table 1. Effect of CRISPRi knock-down of SLLP gene on sporulation in *C. difficile* and *R. lituseburensis*.**

| Target | Spores (%) | | Fold reduction |
|---|---|---|---|
| ***C. difficile* R20291** | **No xylose** | **1% xylose** | |
| negative control | 0.667 | 1.071 | 0.6 |
| CRISPRi::slpA | 1.500 | 0.005 | 362 |
| ***R. lituseburensis* DSM797** | **No xylose** | **1% xylose** | |
| negative control | 28.000 | 57.143 | 0.5 |
| CRISPRi::*sllp* | 11.667 | 0.009 | 1267 |

Data are average of two independent experiments.

**Supplementary Table 2. Bacterial strains used in this study.**

| Species | Strain | Source | Growth medium |
|---|---|---|---|
| *Clostridioides difficile* | R20291[2] | Dr Robert Fagan | TY, BHIS, CDMM |
| *Clostridioides difficile* | FM2.5[2] | Dr Robert Fagan | TY, BHIS, CDMM |
| *Clostridioides difficile* | OPT_2472[3] | Dr Kate Dingle; Optimer Fidaxomicin clinical trial | TY, BHIS |
| *Clostridioides difficile* | CD630 | Dr Trevor Lawley | TY, BHIS |
| *Clostridioides difficile* | OX247Δorf2[1] | Prof. Neil Fairweather (original)<br>Dr Robert Fagan (acquired from) | TY, BHIS |
| *Clostridioides difficile* | OX247[4] | Dr Kate Dingle; Oxford clinical isolates | TY, BHIS |
| *Clostridioides mangenotii* | LM2[5] | Dr William Kelly | PYGS |
| *Clostridioides mangenotii* | G10-CCK1R4-PYG-100 | DSM29133 | DSMZ medium 78 |
| *Asaccharospora irregularis* | VPI 4428 | DSM2635 | DSMZ medium 78 |
| *Acetoanaerobium sticklandii* | HF | DSM519 | DSMZ medium 38 |
| *Romboutsia hominis* | FRIFI | DSM28814 | DSMZ medium 104B |
| *Peptostreptococcus anaerobius* | VPI4330 | DSM2949 | DSMZ medium 78 |
| *Romboutsia ilealis* | CRIB | DSM25109 | DSMZ medium 110A |
| *Romboutsia lituseburensis* | VPI 2751 | DSM797 | DSMZ medium 104B |
| *Paeniclostridium sordellii* | ATCC9714 | DSM2141 | DSMZ medium 78 |
| *Intestinibacter bartlettii* | WAL 16138 | DSM16795 | DSMZ medium 110A |
| *Terrisporobacter glycolicus* | ATCC14880 | DSM1288 | DSMZ medium 110 |
| *Peptoclostridium acidaminophilum* | al2 | DSM3953 | DSMZ medium 454 |
| *Peptoanaerobacter stomatis* | ACC19a | DSM28705 | DSMZ medium 104 |

**Supplementary Table 3. Oligonucleotides used in this study.**

| ID | Sequence | Description |
|---|---|---|
| pIA33_SLLP_2_Rlitu_F | ctataattaaactgtaaatggccaGAATCTACCTTTGCACTTCCgttttagagctagaaatagcaag | CRISPRi, sgRNA in capital letters |
| pIA33_Insert_R | gagaccggtcagatctgcac | CRISPRi |
| pIA33_SlpA1_F | ctataattaaactgtaaatggccaATATCTTCTGCTGCAAACACgttttagagctagaaatagcaag | CRISPRi, sgRNA in capital letters |
| qCdR20291_SlpA_F | AGCTGATGAAGTAGGTCTTGATAAT | RT-qPCR |
| qCdR20291_SlpA_R | TGAGATGCAACTGGAGCTATAC | RT-qPCR |
| qCdR20291_RpoC_F | GCGTGGTCTTATGGCAAATG | RT-qPCR |
| qCdR20291_RpoC_R | GGCACCATGTGAAGATGTAAAG | RT-qPCR |
| qCdR20291_GyrA_F | CGTGCTCTTCCTGATGTTAGAG | RT-qPCR |
| qCdR20291_GyrA_R | GAGCTGACTTCCTGTATGGTTT | RT-qPCR |
| qRlitu_SLLP_F | ACTAAAGCAGAAGCTCCTCAAA | RT-qPCR |
| qRlitu_SLLP_R | CCTTCTACTGTAACTCTTCCTTCTG | RT-qPCR |
| qRlitu_RpoC_F | TAGTGCCAGAAACAGGTGAAA | RT-qPCR |
| qRlitu_RpoC_R | GTTAAAGGATCTCCGGCTACAA | RT-qPCR |
| qRlitu_GyrA_F | AAGACATGCTGAAGGTGAGATAG | RT-qPCR |
| qRlitu_GyrA_R | ATCCGCTGGAACTCTCTTTATG | RT-qPCR |

All primers were synthesised by IDT.

**Supplementary Table 4. X-ray data collection and refinement statistics.**

| | **SlpA H/L Opt2472 (SlpA$_{SLCT2}$)** | **SlpA H/L Ox247 Δorf2 (SlpA$_{SLCT11}$)** |
|---|---|---|
| **Data collection** | | |
| Space group | P1 | P1 |
| Cell dimensions | | |
| *a*, *b*, *c* (Å) | 75.3, 76.5, 78.2 | 47.6, 73.4, 90.8 |
| α, β, γ (°) | 81.8, 78.9, 66.8 | 91.6, 95.7, 96.1 |
| Resolution (Å) | 35.78 - 3.07 (3.18 - 3.07) | 39.08 - 3.10 (3.2 - 3.10) |
| $R_{merge}$ | 0.4827 (1.171) | 0.3859 (1.175) |
| $I / \sigma I$ | 6.2 (3.4) | 7.5 (4.3) |
| Completeness (%) | 99 (99) | 97 (93) |
| Redundancy | 6.6 (6.8) | 6.7 (6.6) |
| *CC1/2* | 0.820 (0.417) | 0.407 (0.159) |
| **Refinement** | | |
| Resolution (Å) | 35.78 - 3.07 (3.18 - 3.07) | 39.08 - 3.10 (3.21 - 3.10) |
| No. reflections | 29118 (2910) | 21675 (2092) |
| $R_{work}$ / $R_{free}$ | 23.1/28.3 (26.1/35.9) | 17.8/23.4 (22.3/28.5) |
| No. atoms | | |
| Protein | 10585 | 8618 |
| Water | 53 | 17 |
| *B*-factors | | |
| Protein | 21.7 | 34.8 |
| Water | 9.1 | 17.5 |
| R.m.s. deviations | | |
| Bond lengths (Å) | 0.003 | 0.003 |
| Bond angles (°) | 0.5 | 0.59 |
| PDB ID | 9F8F | 9F8E |

Data collected from a single crystal for each SlpA

*Values in parentheses are for highest-resolution shell.

**Supplementary Data Inventory**

**SupplementaryData1**

Referenced table of representative species with clear evidence of S-layer including those with robustly verified SLLP presented in Fig.1, including identified available experimental evidence, available structures, previously verified SLLP protein sequences, sequences of SLLPs identified in this study, and no-AD sequences used in Supplementary Fig. 1b and c.

**SupplementaryData2**

*Clostridioides difficile, Bacillus, Peptostreptococcaceae*, and Firmicutes genomes used in bioinformatic analyses.

**SupplementaryData3**

Lists of orthologues used for the inference of phylogenetic species trees.

**SupplementaryData4**

Species phylogeny of 453 *Peptostreptococcaceae* from a concatenated alignment of 111 orthologue protein sequences.

**SupplementaryData5**

Species phylogeny of 828 Firmicutes from a concatenated alignment of 84 orthologue protein sequences.

**SupplementaryData6**

Species phylogeny of 4,774 *Bacillus* from a concatenated alignment of 84 orthologue protein sequences.

**SupplementaryData7**

Protein sequences of the identified *Bacillus* SAP/EA1/SAP2 and *C. difficile* SlpA homologues, and gene sequences of unique *C. difficile slpA*.

**SupplementaryData8**

Full-length and AD-removed protein sequences of cell wall anchored protein homologues presented in Supplementary Fig. 3.

**SupplementaryData9**

Full-length protein sequences of cell wall anchored protein homologues presented in Fig. 2.

**SupplementaryData10**

Full-length and AD-removed protein sequences of cell wall anchored protein homologues presented in Fig. 3a-c, and no-AD proteins identified as co-distributing with SLH or CWB2 anchoring systems presented in Fig. 3f.

**SupplementaryData11**

*C. difficile* core genome species tree generated with IQ-TREE[6] under LG + G4 model

**SupplementaryData12**

OrthoFinder analysis of 266 *C. difficile* genomes.

**SupplementaryData13**

*C. difficile* SlpA positive/negative selection analyses.

**SupplementaryData14**

PDB validation report for 9F8E $SlpA_{SLCT11}$ crystal structure.

**SupplementaryData15**

PDB validation report for 9F8F $SlpA_{SLCT2}$ crystal structure.
